# Niclosamide inhibits SARS-CoV2 entry by blocking internalization through pH-dependent CLIC/GEEC endocytic pathway

**DOI:** 10.1101/2020.12.16.422529

**Authors:** Chaitra Prabhakara, Rashmi Godbole, Parijat Sil, Sowmya Jahnavi, Thomas S van Zanten, Dhruv Sheth, Neeraja Subhash, Anchal Chandra, Vijay Kumar Nuthakki, Theja Parassini Puthiyapurayil, Riyaz Ahmed, Ashaq Hussain Najar, Sai Manoz Lingamallu, Snigdhadev Das, Bhagyashri Mahajan, Praveen Vemula, Sandip B Bharate, Parvinder Pal Singh, Ram Vishwakarma, Arjun Guha, Varadharajan Sundaramurthy, Satyajit Mayor

## Abstract

Many viruses utilize the host endo-lysosomal network to infect cells. Tracing the endocytic itinerary of SARS-CoV2 can provide insights into viral trafficking and aid in designing new therapeutic targets. Here, we demonstrate that the receptor binding domain (RBD) of SARS-CoV2 is internalized via the clathrin and dynamin-independent, pH-dependent CLIC/GEEC (CG) endocytic pathway. Endosomal acidification inhibitors like BafilomycinA1 and NH_4_Cl, which inhibit the CG pathway, strongly block the uptake of RBD. Using transduction assays with SARS-CoV2 Spike-pseudovirus, we confirmed that these acidification inhibitors also impede viral infection. By contrast, Chloroquine neither affects RBD uptake nor extensively alters the endosomal pH, yet attenuates Spike-pseudovirus entry, indicating a pH-independent mechanism of intervention. We screened a subset of FDA-approved acidification inhibitors and found Niclosamide to be a potential SARS-CoV2 entry inhibitor. Niclosamide, thus, could provide broader applicability in subverting infection of similar category viruses entering host cells via this pH-dependent endocytic pathway.

## Introduction

Coronaviruses (CoVs) are a group of related enveloped RNA viruses of which two alpha CoVs (229E and NL63) and four beta CoVs (OC43, HKU, SARS, and MERS) are known to cause respiratory tract infections in humans. The recent emergence of SARS-CoV2 and its rapid spread across the world has posed a global health emergency ^1^. While several therapeutic strategies are currently being used to alleviate the respiratory symptoms of patients infected with SARS-CoV2 ^2,3^, there is limited understanding of the cell biology of viral entry as well as the availability of drugs which target this process. A search for antivirals affecting the endocytic entry of viruses is particularly exciting as infections from multiple related viruses can be controlled through the inhibition of a common step.

Virus entry into host cells is a multistep process. A key step in successful invasion is the release of viral genomic content into the host cell cytoplasm. To achieve this, viruses bind to specific cell surface receptors and subsequently undergo membrane fusion either directly at the plasma membrane or following endocytic uptake. While fusion directly at the plasma membrane is well established for HIV and Influenza virus infections ^4,5^, both alternatives of entry are feasible for CoV infections depending on the availability of receptors and proteases at the host cell surface. Different CoVs interact with a range of specific receptors for entry. For instance, CoV 229E binds to CD13 (aminopeptidase N) ^6^, CoVs OC43 and HKU1 recognize 9-O-acetylated sialic acids ^7^, MERS uses DPP4/CD26 ^8^ and CoVs NL63 ^9^, SARS-CoV ^10^ and SARS-CoV2 ^11^ interact with <u>a</u>ngiotensin <u>c</u>onverting <u>e</u>nzyme 2 (ACE2). Although ACE2 is a well studied receptor, other receptors for SARS-CoV2 are being discovered ^12–16^. Additionally, CoVs require proteolytic processing of the viral envelope spike protein by host cell proteases to gain entry ^17,18^. Therefore, these viruses can directly fuse at the cell surface if the Spike protein is cleaved by a cell surface serine protease like TMPRSS2 ^11,19^, or utilize an endo-lysosomal route for fusion, where the Spike protein is primed by cysteine protease Cathepsins ^11,20–22^.

The role of the endo-lysosomal network appears to be crucial in delivering these viruses to acidic compartments. Cathepsins function optimally in a low pH environment ^17,23^. Inhibitors of acidification which increase the pH of endosomal compartments significantly reduce the infection of spike pseudotyped as well as native MERS-CoV, SARS-CoV, SARS-CoV2 viruses ^11,24–27^. Supporting this view, drugs inhibiting the maturation of late endosome to lysosome, Apilimod and YM201636, also reduce MERS, SARS-CoV, SARS-Cov2 infection ^24,28,29^. These studies emphasize the importance of a pH-dependent endocytic route in viral entry and infection. However, the endocytic routes preferred by SARS-CoV2 for host cell entry are largely unknown.

Multiple endocytic pathways operate at the cell surface ^30,31^. One of these, the clathrin and dynamin independent CLIC/GEEC (CG) endocytic pathway ^32^, is of particular interest here as uptake through this pathway is known to be pH-dependent. Vacuolar ATPases (V-ATPases), which actively pump protons into the endocytic compartments ^33^, play a crucial role in the formation of CG endosomes as established using genetic and pharmacological perturbations ^34,35^. By contrast, uptake through <u>c</u>lathrin <u>m</u>ediated <u>e</u>ndocytosis (CME) remains unaltered upon V-ATPase perturbation ^34^. The homotypic fusion of nascent CG endosomes (called CLICs – <u>cl</u>athrin-independent <u>c</u>arriers) forms highly acidic early endosomal compartments of the CG pathway (called GEECs – <u>G</u>PI anchored protein <u>e</u>nriched <u>e</u>arly endosomal <u>c</u>ompartments) with an estimated luminal pH of 6.0 ^36^. Thus, GEECs could provide a conducive environment for viral uncoating and membrane fusion. Additionally, V-ATPases and ARP2/3 complex, both imperative for CG endocytosis ^37^, are identified as host factors necessary for SARS-CoV2 viral infection in genome-wide loss of function screen ^38^. Interestingly, Adeno-associated virus (AAV2) hijacks the CG pathway for infection ^39^ and SARS-CoV has also been reported to enter cells through a clathrin and dynamin independent endocytic pathway ^26^. These observations prompted us to study the role of CG endocytosis in the context of SARS-CoV2 entry and infection.

In this report, we show that the receptor <u>b</u>inding <u>d</u>omain (RBD) of SARS-CoV2 Spike protein is endocytosed through the CG pathway and its uptake is sensitive to pharmacological perturbations of the CG pathway. RBD uptake, similar to CG cargo uptake, is strongly inhibited by acidification inhibitors such as BafilomycinA1 and NH_4_Cl. Inhibitors of endosomal acidification also blocked infection by SARS-CoV2 Spike-pseudoviruses. Extending our observations, we conducted a targeted screen using a subset of FDA-approved drugs which are known to interfere with endosomal acidification. We identified Niclosamide as a promising candidate that inhibits RBD uptake, Spike-pseudovirus infection and, in combination, potentiates the effects of Hydroxychloroquine. We suggest that Niclosamide could be a feasible start point for developing small molecule entry inhibitors to mitigate SARS-CoV2 infection.

## Results

### Generation of SARS-CoV2 probe to study its endocytosis itinerary

The Spike (S) protein of SARS-CoV2 plays crucial roles in mediating viral entry to cells. S protein binds to the receptors on the host cell surface through the S1 subunit which harbours the receptor binding domain (RBD) and aids in membrane fusion through the S2 subunit ^40^. To explore the trafficking route of SARS-CoV2 in human cells, we purified RBD protein, following a previously established method ^41^, and generated fluorescently labelled RBD using NHS-ester chemistry (Figure S1A, S1B, Methods). We chose human adenocarcinoma gastric cells (AGS cells) as a model system to study RBD uptake as the cell line exhibits multiple endocytic routes ^42^ and is also permissive to infection by SARS-CoV2 Spike-pseudovirus (Figure S6E). We tested the specificity of the labelled RBD probe in AGS cells transiently overexpressing myc-tagged ACE2 and found that more RBD was bound to cells overexpressing ACE2 (Figure S1C). We observed a positive correlation between the amount of RBD endocytosed and surface levels of ACE2 (Figure S1D, S1E), supporting the notion that ACE2 is one of the cell surface receptors of RBD ^11^.

### RBD is internalized via CG endocytosis and RBD uptake is sensitive to CG Pathway inhibitors

We employed the methodology of tracking RBD uptake along with cargoes specific to CME (transferrin) and CG (10kDa dextran) endocytic pathway to determine the endocytic route taken up by RBD (Figure 1A). AGS cells without overexpression of ACE2 also support RBD uptake suggesting that there is sufficient endogenous receptor expressed in these cells. At 10 minutes post internalization, transferrin endosomes of the CME pathway are distinct from dextran endosomes of the CG pathway. At these times, internalized RBD is colocalized with endosomes containing the CG cargo but not the CME cargo (Figure S2A, S2B). At 30 minutes post internalization, as well, this segregation remains. While a small fraction of RBD endosomes were colocalized with endosomes containing both transferrin and dextran, a large fraction of RBD endosomes were localized to compartments uniquely marked by dextran (Figure 1B, 1C; compare % RBD with transferrin and dextran). This suggests that the itinerary of uptake of RBD is similar to CG cargo and different from CME cargo.

**Figure 1:**
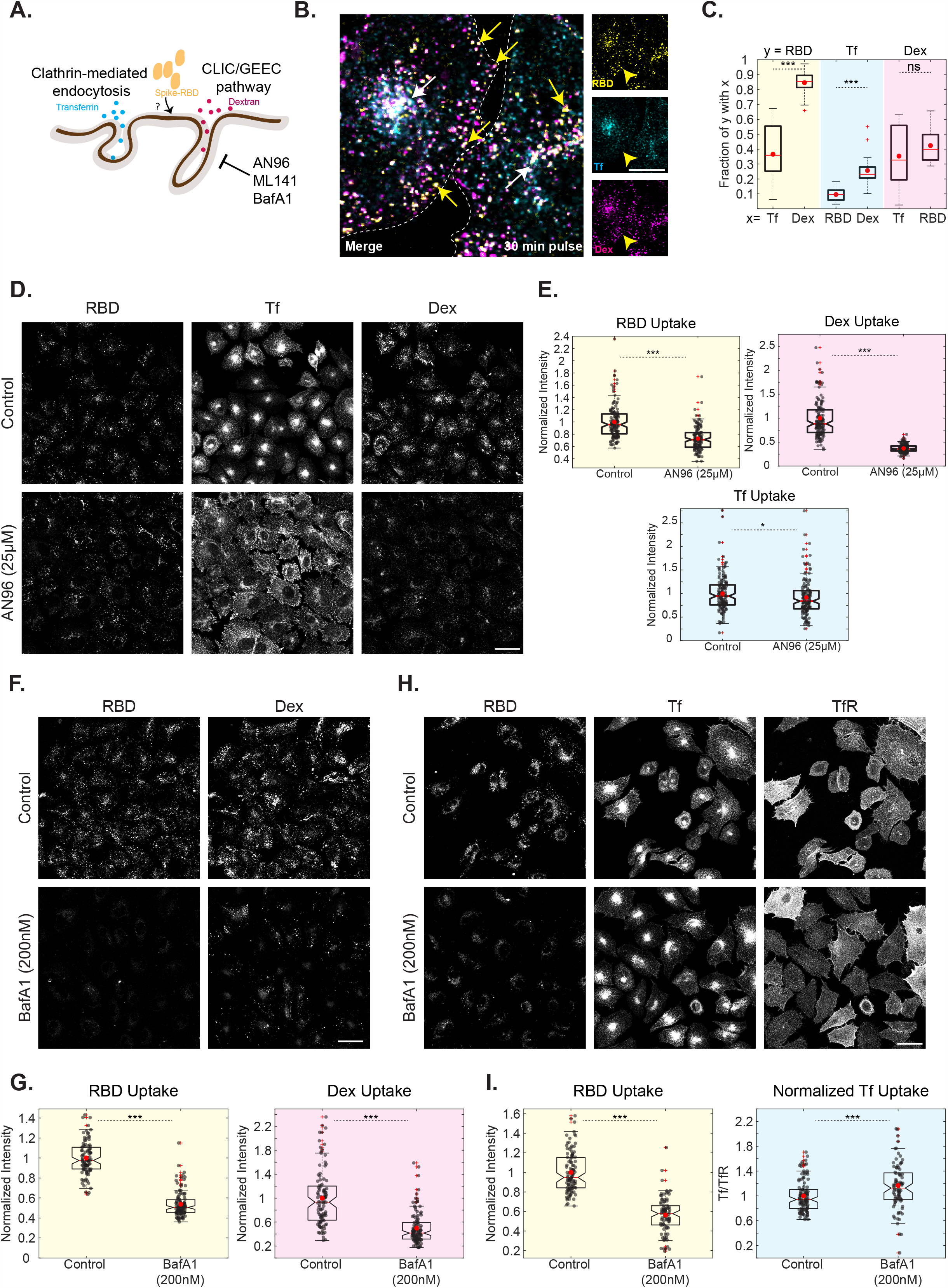
RBD uptake is sensitive to CG Pathway inhibitors in AGS cells. A: Schematic describing endocytic pathways at the plasma membrane with specific cargoes for each endocytic pathway: transferrin (CME Cargo) and 10kDa dextran (CG Cargo). AN96, ML141 and BafA1 specifically affect the uptake of CG cargoes. B, C: AGS cells were pulsed with RBD, dextran and transferrin for 30 minutes and imaged at high resolution after fixation. Images are shown in B and quantification of Manders’ co-occurrence coefficient is shown in C. This compares the fraction of RBD endosomal intensity with transferrin or dextran (p-value < e-06), transferrin endosomal intensity with dextran or RBD (p-value < e-05) and dextran endosomal intensity with transferrin or RBD (p-value = 0.18). RBD is more co-localized to dextran endosomes. Number of cells = 10. White arrow represents endosomes containing RBD, dextran and transferrin. Yellow arrow represents endosomes with RBD and dextran without transferrin. Dashed white line in B represents the approximate cell boundary. D, E: AGS cells were pretreated with Control (0.6% DMSO) or AN96 25µM for 30 minutes and pulsed with RBD, dextran and transferrin for 30 minutes with or without the inhibitor. Treatment with AN96 reduces RBD (p-value < e-19) and dextran (p < e-44) uptake while minimally alters transferrin uptake (p = 0.02). Images are shown in D and quantification in E. Numbers of cells > 100 for each treatment. F, G: AGS cells were treated with Control (0.2% DMSO) or BafA1 200nM for 30 minutes and pulsed with RBD and dextran for 30 minutes with or without the inhibitor. Treatment with BafA1 reduces RBD (p-value < e-33) and dextran (p-value < e-18) uptake. Images are shown in F and quantification in G. Numbers of cells > 100 for each treatment. H, I: AGS cells were treated with Control (0.2% DMSO) or BafA1 200nM for 30 minutes and pulsed with RBD and transferrin for 30 minutes with or without the inhibitor. The surface transferrin receptor (TfR) was labelled after fixation. Treatment with BafA1 reduces RBD uptake (p-value < e-27) and increased normalized transferrin uptake (p-value < e-03). Images are shown in H and quantification in I. Numbers of cells > 80 for each treatment. Data (E, G, I) is represented as a scatter with box plot. Black dots represent per-cell data points. Box plot represents the distribution (25% to 75% percentile) with the red line indicating the median and red dot indicating the mean of the distribution. Whiskers represent distribution up to 1.5 times interquartile range and + indicates outliers beyond the whiskers. In the entire manuscript, ***, **, * and ns indicate p-value of Wilcoxon rank-sum test < 0.001, <0.01, <0.05 and not significant, respectively. Scale bar: 20µm (B) and 40µm (D, F, H).

Towards determining the trafficking route of RBD, we examined the effect of inhibitors of CG pathway on RBD, dextran and transferrin uptake in AGS cells. Cells were subjected to a brief pre-treatment with different inhibitors (30 minutes), followed by a pulse of RBD, dextran and transferrin (30 minutes) in the presence of these inhibitors (Methods). CG pathway is regulated by small GTPases - CDC42 ^43^, Arf1 ^44^ and GEF of Arf1, GBF1 ^45^. Inhibitors that block the function of these regulators affect the formation of CG endosomes without altering uptake through the CME pathway. The inhibitor AN96, which is a stable analog of LG-186 ^46,47^, targets GBF1 and specifically affects the CG pathway (Godbole et al., Manuscript in preparation). We observed that AN96 treatment reduced both RBD and dextran uptake but had minimal effects on the amount of transferrin internalized (Figure 1D, 1E). We also observed that the peri-nuclear transferrin recycling endosomal pool was redistributed throughout the cytoplasm upon treatment with AN96 without affecting the net amount of transferrin internalized. Another CG pathway inhibitor, ML141 (CDC42 inhibitor) ^47^, also significantly decreased both dextran as well as RBD uptake (Figure S2E, S2F).

Blocking of the CG pathway often results in the redistribution of CG cargo towards CME pathway ^42^. Therefore, using high-resolution imaging, we assessed the fate of the RBD endosomes that continued to be internalized upon treatment with AN96. We observed that an increased fraction of internalized RBD endosomes colocalized with transferrin in AN96 treated cells compared to that of control (Figure S2C, S2D(i)). Similarly, increased co-occurrence was observed in the fraction of dextran endosomes associated with transferrin endosomes on comparing the control with AN96 treated cells (Figure S2D(iii)). Blocking the CG pathway results in altered trafficking itinerary of RBD and increases its association with transferrin. This suggests that RBD could be redirected to be internalized via the CME upon blocking the CG pathway.

Since 10kDa dextran marks both CG cargo as well as larger endocytic compartments like those derived from macropinocytosis ^48^, we tested if macropinocytosis plays any role in RBD uptake. Several viruses utilize macropinocytosis pathway as an entry route into cells ^49^. Macropinocytosis is dependent on amiloride-sensitive Na+/H+ exchangers ^50^. Upon treatment with Amiloride, we found no alteration in the uptake of RBD, dextran and transferrin confirming that macropinocytosis does not play a role in RBD trafficking in AGS cells (Figure S2G, S2H). Together, the co-localization studies and pharmacological inhibition experiments strongly suggest that RBD uptake occurs via the CG Pathway and is inhibited by specific blockers of the CG pathway.

### RBD and CG uptake is blocked by endosomal acidification inhibitors – BafilomycinA1 and NH_*4*_*Cl*

Given the relevance of acidification in both formation of CG endosomes ^34,36^ and in the context of viral infection ^51^, we focused on studying the role of acidification inhibitors on the uptake of RBD. We checked the effect of BafilomycinA1 (BafA1), a specific inhibitor of V-ATPase ^52^, on RBD, dextran and transferrin uptake in AGS cells. Treatment with 200nM BafA1 strongly reduced both RBD and dextran uptake (Figure 1F, 1G) and enhanced the normalized transferrin uptake (Figure 1H, 1I). This could be because BafA1 also retards the transferrin recycling from the recycling endosomes ^53^ and thereby increasing the net amount of transferrin internalized within cells as observed. A dose-dependent reduction in RBD and dextran uptake and an increase in transferrin uptake was seen when cells were treated with a higher concentration of BafA1 (Figure S3A, S3B (i)). We examined the effect of NH_4_Cl, a weak base known to alter endosomal acidification ^54^, on the uptake of these 3 cargoes. We observed similar results as with BafA1 (Figure S3A, S3B (i)), thus re-establishing our earlier ^34^ finding that uptake via the CG pathway is pH sensitive and blocking acidification results in reduced CG uptake.

Towards understanding the mechanism of action for acidification inhibitors in bringing about these changes in trafficking, we assessed their effect on two parameters – numbers of endosomes (Figure S3B (ii)) and per-endosome intensity in the presence/absence of inhibitor (Figure S3B (iii)). We observed that both BafA1 and NH_4_Cl reduced the total number of RBD and dextran endosomes without affecting the per-endosome intensity. However, while the total number of transferrin endosomes remained unchanged, the per-endosome intensity of transferrin increased with BafA1 and NH_4_Cl treatment. This indicates that the reduction in RBD and dextran is likely due to a block in the entry while an increase in per-endosome transferrin intensity could be because of a block in the formation of recycling endosome carriers, as proposed earlier.

We studied the effect of BafA1 on RBD uptake in cells overexpressing myc-tagged ACE2 receptor by measuring the uptake of RBD normalized to the surface ACE2 levels. We observed that BafA1 strongly affects the normalized RBD uptake (Figure S3C, S3D). HEK-293T cells, which is also permissive to Spike-pseudovirus transduction (Figure S6C), showed similar inhibition of RBD and dextran uptake, and increase in transferrin uptake with BafA1 (Figure S10A-S10D).

### RBD is localized to acidic compartments

Internalized cargoes can be recycled along with the bulk membrane ^55^ or directed towards degradation with the fluid phase ^56^. Typically, transferrin bound to its receptor marks the early sorting and recycling endosomes and lysotracker labels the acidic degradative compartments within a cell ^31^. At 30 minutes of pulse with RBD, dextran and transferrin, while a small fraction of RBD (∼36%) associated with transferrin, the majority of RBD (∼84%) co-localized with dextran suggesting that RBD is directed predominantly towards the degradation route rather than the recycling route (Figure 1B, 1C). The lysotracker labelling showed highly acidic tubular compartments with significant co-localization with RBD. At 30 minutes of pulse with RBD, around 55% of RBD co-localized with lysotracker marked compartments. At longer time points (3 hours) of pulse with RBD, an even increased proportion of RBD (85%) associated with compartments marked by lysotracker, confirming that RBD is trafficked to acidic compartments (Figure 2A, 2B).

**Figure 2:**
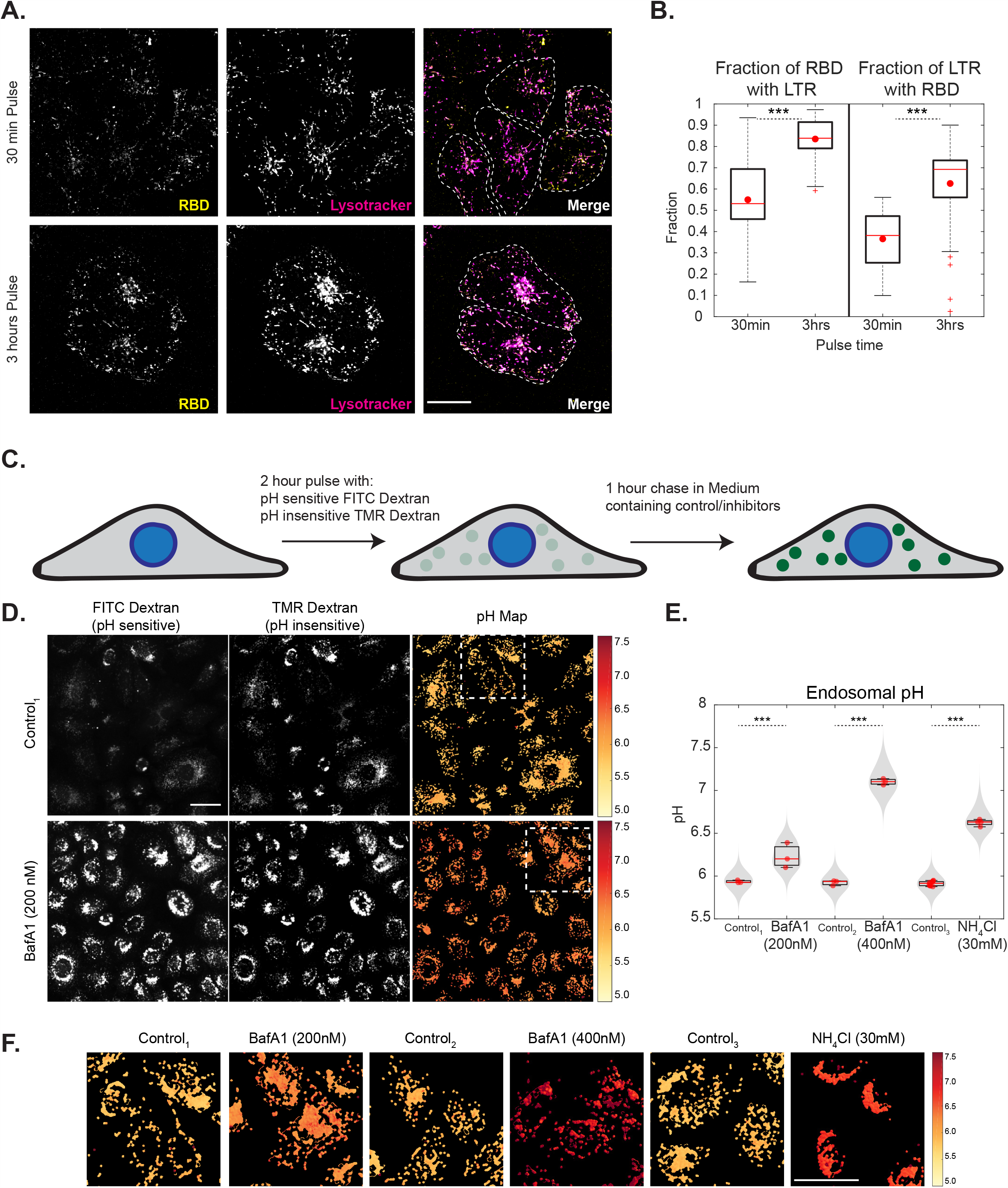
RBD trafficking with CG cargo is localized to acidic compartments and endosomal acidification inhibitors neutralize these endosomes. A, B: AGS cells were pulsed with RBD for 30 mins or 3 hours, labelled with lysotracker in the last 15 minutes of pulse and imaged live at high resolution. Images are shown in A and quantification of Manders’ co-occurrence coefficient is shown in B. RBD is colocalized with Lysotracker positive compartments. With increase in time more RBD is associated with Lysostracker (p-value < e-06) and more Lysotracker positive compartments have RBD (p-value < e-06). Each condition has >12 cells. Dashed white line in A represents approximate cell boundary. C: Schematic describing the experimental protocol for estimating the pH of endosomes by ratiometric measurements using pH-sensitive (FITC) and pH-insensitive (TMR) dextran. D-F: AGS cells were pulsed with FITC and TMR dextran for 2 hours, chased for 1 hour with BafA1 200nM/400nM, NH_4_Cl 30mM or control and imaged live. Endosomal pH is increased upon addition of acidification inhibitors (p-values < e-118 for BafA1 200nM, < e-122 for BafA1 400nM, < e-223 for NH_4_Cl). Images along with pH maps are shown in D (and in S4C) and quantification in E (and in S4D). Enlarged regions of pH maps indicated by white boxes are shown in F. Box plot in E represents the distribution of medians of each repeat which is denoted by red dots. Violin plot indicates all the data points from repeats. Colour bar in F corresponds to the estimated endosomal pH. Control_1_ is 0.2% DMSO, Control_2_ is 0.4% DMSO and Control_3_ is 0% DMSO. Number of repeats ≥ 3 for each treatment and each repeat has >80 cells. Scale bar: 20µm (A) and 40µm (D).

### BafilomycinA1 and NH_4_Cl alter the pH of acidic endosomal compartments

We next focused on determining the change in endosomal pH brought about by various inhibitors within the acidic compartments populated by RBD. Cells were labelled with pH-sensitive (FITC) and pH-insensitive (TMR) dextran for 2 hours and chased for 1 hour with or without inhibitors (Figure 2C, Methods). The above pulse and chase durations were chosen to allow accumulation of labelled dextran in late endosomes and lysosomal compartments (co-labelled by Lysotracker, data not shown). Additionally, since the acidification inhibitors also have inhibitory roles in the early steps of CG endocytosis as discussed in the previous section, to evaluate their effect on endosomal pH, cells were incubated with inhibitors only during the chase. While the ratio of the fluorescence of these probes is used to estimate endosomal pH by comparing the ratio with the calibration curve ^57^ (Figure S4A, S4B, Methods), quantifications of the endosomal intensities and the endosomal number of TMR dextran aids in understanding the effect of various drugs on late endosomal trafficking.

Treatment of cells with acidification inhibitors showed an increase in endosomal pH. The average pH of the late endosomes in control cells was 5.8. The pH of these compartments increased to 6.2 and 7.1 in the presence of BafA1 200nM and 400nM respectively. Incubation with NH_4_Cl also resulted in increasing the pH of these endosomes to 6.6 (Figure 2D, 2E, S4C). While BafA1 marginally changed the TMR intensity per endosome, NH_4_Cl greatly increased the TMR intensity indicating that NH_4_Cl also brings about the fusion of endosomes (Figure S4D). All the acidification inhibitors also reduced the numbers of endosomes (Figure S4D) and this effect was most prominent with NH_4_Cl wherein the endosomes were organized close to the perinuclear region (Figure S4C). The spatial pH maps show the distribution of pH of endosomes within a cell. Cells treated with BafA1 400nM and NH_4_Cl showed a homogenous distribution of endosomes with increased pH similar to the respective cell averages. BafA1 200nM cells, on the other hand, showed heterogeneity in endosomal pH with some endosomes depicting high pH while others were closer to the average (Figure 2F).

To assess the effect of BafA1 on the pH of early time point endosomes, AGS cells were labelled with FITC and TMR dextran for 20 minutes and chased for 10 minutes with or without BafA1 for the entire duration of pulse and chase (Figure S4E). While the total amount of dextran uptake is not affected significantly, the endosomal FITC intensity and the endosomal ratio of FITC/TMR, which can be considered as a proxy for endosomal pH, show a robust increase with BafA1 treatment (Figure S4F). This indicates that BafA1 also affects the endosomal pH of early time point endosomes.

### Chloroquine treatment does not affect RBD uptake and minimally alters endosomal pH

Chloroquine, a diprotic weak base, is expected to accumulate in acidic compartments and neutralize lysosomal pH ^58^. While, mounting evidence shows that Chloroquine and its analogs can inhibit the infection by several viruses such as Ebola, Dengue, Chikungunya, HIV, etc ^59^, many studies point towards differences between the mode of action of Chloroquine and acidification inhibitors – BafA1 and NH_4_Cl ^60,61^. We, therefore, tested the effect of Chloroquine on RBD, dextran and transferrin uptake to verify if it behaves like BafA1. We found that upon treatment with Chloroquine, the uptake of neither RBD nor transferrin was altered significantly (Figure S5A, S5B). Dextran uptake was marginally higher upon treatment with Chloroquine (Figure S5B).

The effect of Chloroquine in changing the endosomal pH of late endosomes using FITC/TMR ratio as a proxy for endosomal pH was assessed. At different concentrations of Chloroquine tested, the endosomal pH was only minimally increased (Figure S5C, S5D). We also observed that both FITC and TMR endosomal intensities increased with the concentration of Chloroquine. To confirm our results, we used another method to estimate endosomal pH. FITC has a pH-sensitive (488nm) and a pH-insensitive excitation (450nm) ^54^. We used the 488/458 excitation ratio of FITC dextran as a readout of pH and found that this ratio also showed only a small albeit significant increase with Chloroquine when compared to control cells, unlike the increase brought about by NH_4_Cl (Figure S5E, S5F). This observation could explain the lack of an endocytic effect on RBD, dextran or transferrin uptake upon treatment with Chloroquine.

### Designing Spike-pseudovirus transduction assay to specifically address the effects of inhibitors on viral entry

To ascertain that the observations made using the RBD, as a proxy for viral entry, are valid in the context of a viral entity decorated by the SARS-CoV2 Spike protein itself, we generated SARS-CoV2 Spike-pseudotyped lentiviral particles (Spike-pseudovirus), following a previously established methodology ^62^. mCherry fluorescent protein expression was used as a reporter for assessing viral infection (Figure S6A, Methods). The expression of the Spike protein in the pseudovirus particles was verified by a western blot using the antibody against the C-terminal Strep-tag of Spike, which revealed bands corresponding to both the S2 fragment as well as the full-length protein (Figure S6B). The infection specificity of the pseudovirus was validated by infection of human (HEK-293T) versus mouse (NIH-3T3) cells, where the latter showed lower infectivity, consistent with a lack of a bonafide hACE2 receptor to bind the Spike protein (Figure S6C). Independently, a competition experiment was conducted to check the effect of excess soluble RBD on transduction of Spike-pseudovirus in HEK-293T (Methods). The transduction efficiency was reduced in the presence of soluble RBD, indicating that the Spike-pseudovirus competes for the same binding sites as RBD (Figure S6D). However, the inhibition was not complete, possibly since even high concentration of free RBD in the solution cannot compete with the high effective concentrations on Spike-pseudoviruses, augmented even more by their trimeric configuration that facilitates multi-valent interactions ^63^.

Since our experiments were aimed at understanding the entry mechanism of Spike-pseudovirus, we designed the transduction assays with shorter pseudovirus incubation time and followed the infection efficiency by tracing reporter gene expression at a later time point. We characterized the transduction efficiency of the pseudovirus as a function of its MOI and time of incubation to obtain an optimum MOI and incubation time (Figure S6E). Transduction efficiency measured across the tested regime suggested 4 or 8 hours of incubation at 0.5 MOI to be optimal to achieve at least >1000 positive cells (per well of a 96-well assay plate) with reasonably low viral load and incubation time.

### Spike-pseudovirus transduction is reduced by endosomal acidification inhibitors and Chloroquine

If the pseudovirus expressing the full-length spike mimicked the same trafficking pathway for entry as RBD, we reasoned that the transduction efficiency would be reduced upon treatment with inhibitors affecting RBD uptake. SARS-CoV2 Spike-pseudovirus transduction efficiency has been reported to be reduced upon treatment with NH_4_Cl, BafA1 and Chloroquine ^24,64^. However, we wanted to specifically explore the actions of these inhibitors at the initial stages of infection. Therefore, to address this, we pre-treated cells with drugs for an hour followed by the addition of Spike-pseudovirus in the presence of the drug for 2, 4 or 8 hours. Both virus and drug were removed thereafter, and cells were incubated with fresh media either in the absence or presence of a minimal concentration of the drug as indicated (Figure 3A, Methods). This design was chosen to reduce long-term toxicity of the inhibitors to the cells and minimize any secondary effects on the translational processes of the reporter gene post entry. Infection, or transduction efficiency, is reported as the normalized percentage of transduction compared to corresponding control and cell viability is measured in terms of nuclei number normalized to control.

**Figure 3:**
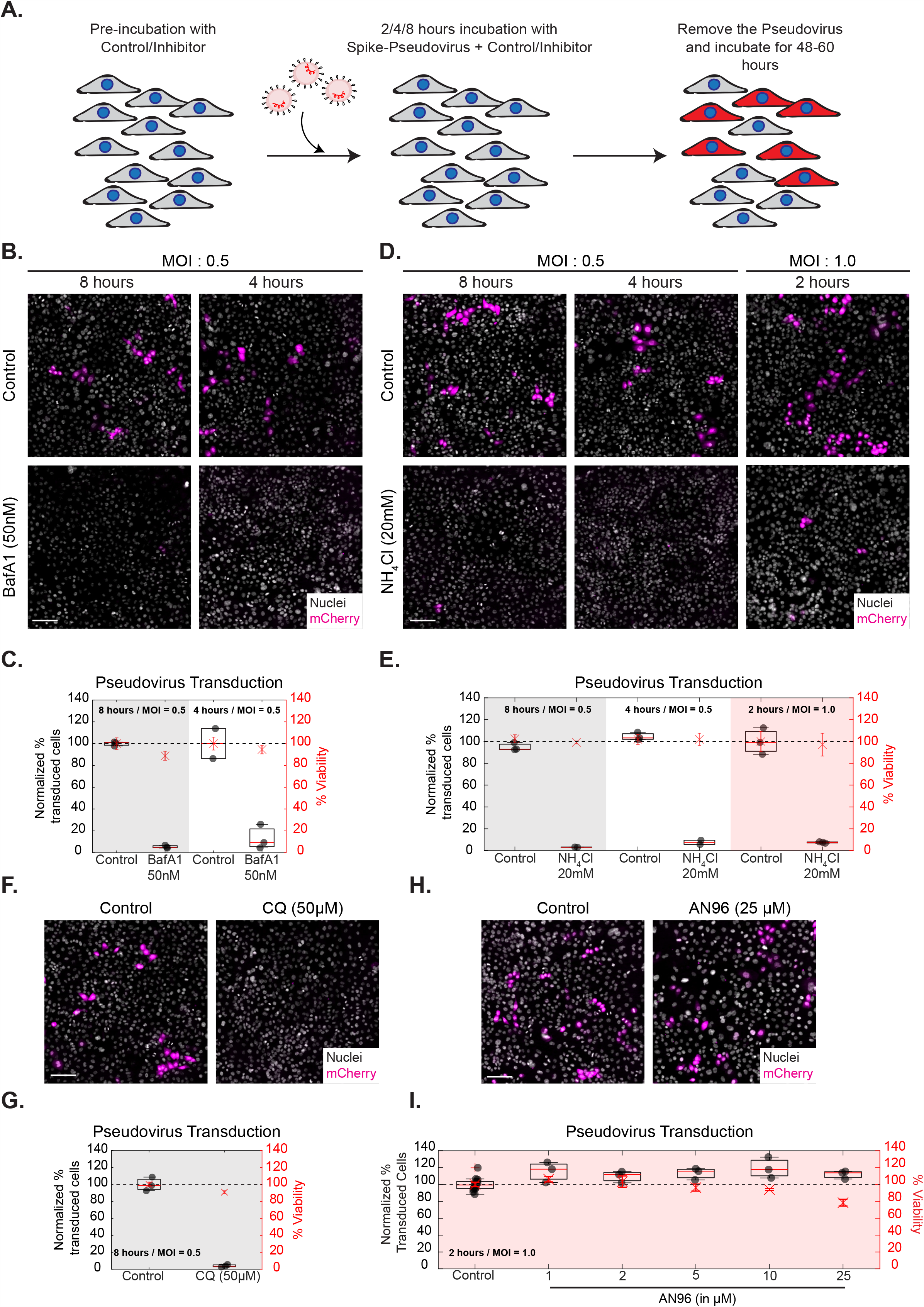
BafA1, NH_4_Cl and Chloroquine affect Spike-pseudovirus infection. A: Schematic describing the experimental protocol for SARS-CoV2 Spike-pseudovirus transduction assay. (B-I): AGS cells were pre-incubated with the inhibitors (BafA1 50nM, NH_4_Cl 20mM, CQ 50µM and AN96 1-25µM) for an hour and transduced with Spike-pseudovirus at MOI 0.5 or MOI 1.0 for indicated incubation times. Following this, virus and inhibitor-containing medium was removed and cells were further incubated with either inhibitor-free media (B-G) or low concentration of the inhibitor-containing media (1µM of AN96 in H-I). B, D, F and H show images of AGS cells expressing the reporter mCherry protein and C, E, G and I show the quantification of normalized % transduction of cells treated with inhibitors compared to respective controls. B-E: Transduction efficiency is reduced with BafA1 (p-values < e-86 for 8 hours, < e-60 for 4 hours) and NH_4_Cl (p-values < e-83 for 8 hours, < e-72 for 4 hours, < e-85 for 2 hours) at all time points of incubation. Control in B is 0.05%DMSO and in D is 0%DMSO. Number of repeats ≥ 2 for each treatment. F, G: Transduction efficiency is reduced with CQ with 8 hours of incubation (p-value < e-90). Number of repeats = 3 each for 0% DMSO Control and CQ. H, I: No reduction in efficiency is seen upon treatment with AN96 with 2 hours of incubation across all the concentrations tested compared to the pooled control from 0.6%, 0.24%, 0.12%, 0.048% and 0.024% DMSO treatments. Number of repeats = 15 for pooled control and 3 for each concentration of AN96. Data (C, E, G, I) is represented as percentage transduction normalized to control on left Y-axis along with percentage viability, calculated from the total number of nuclei, normalized to control on the right Y-axis. Black dots represent the mean of each repeat. Box plot represents the distribution of the means (25% to 75% percentile) with red line indicating the median of the distribution. Asterix with error bars in red represents the mean +/-SD of % viability. Black dotted line marks 100%. Scale bar: 100µm (B, D, F, H).

We tested the effect of NH_4_Cl and BafA1 on the Spike-pseudovirus transduction assay in AGS cells upon treatment with the 20mM NH_4_Cl or 50nM BafA1. At the end of designated time points, the pseudovirus containing media along with the NH_4_Cl and BafA1 was removed and replenished with fresh media alone. We observed a significant reduction of Spike-pseudovirus transduction with NH_4_Cl and BafA1 compared to the corresponding controls with no significant difference in cell viability at the end of both 4 and 8 hours (Figure 3B-3E). NH_4_Cl also shows robust reduction even upon 2 hours of incubation (Figure 3E). HEK-293T cells also exhibited a similar inhibition of transduction upon treatment with 20mM NH_4_Cl or 50nM BafA1 (Figure S10 E-G).

Although Chloroquine did not alter RBD uptake or increase the pH of the endocytic compartments significantly, we observed a marked reduction of viral transduction with 50µM Chloroquine treatment in AGS cells (Figure 3D, 3E) and 10µM Chloroquine treatment in HEK-293T cells (Figure S10E-G), where the drug was removed along with the Spike-pseudovirus at the end of 8 hours of incubation. Upon testing whether long term incubation of Chloroquine (as used in this assay), results in changes in endosomal pH or RBD uptake, we observed no alterations in the assessed phenotypes in AGS cells (Figure S5G, S5H). This suggests a distinct pH-independent mechanism of intervention by Chloroquine, functioning at the initial stages of infection.

RBD uptake is reduced upon treatment with AN96 and ML141, albeit to a lesser extent compared to the effect of BafA1 and NH_4_Cl. Therefore, we assessed the effect of AN96 and ML141 on Spike-pseudovirus transduction in AGS cells, using the 2 hours and 8 hours format of the assay, respectively. The AN96 concentration was reduced to a non-toxic level of 1µM after removal of the virus. With this experimental design, 5 different concentrations of AN-96 were tested, and we observed no reduction in normalized percentage transduction even at the highest concentration of 25µM (Fig. 3F and G), with no compromise on cell viability. This was consistent with the observation that RBD is rerouted and associates more with transferrin. In case of ML141, we treated the cells with 5µM of the drug and observed no difference in normalized percentage transduction compared to the control (Figure S6F, S6G). Partial inhibition of uptake may not strongly manifest in our pseudovirus assay, as the read-out is all or none and is not sensitive to the number of virus particles entering the cells. Our findings suggest that inhibitors that affect both RBD uptake and neutralize acidic endosomes could be one of the strategies used to impede Spike-pseudovirus transduction.

### Identifying FDA-approved drugs functioning similar to BafA1 and NH_*4*_*Cl*

Armed with the knowledge on the mode of action of acidification inhibitors in reducing the uptake of RBD, increasing the pH of endosomes and abrogating the infection of Spike-pseuodovirus, we screened a small subset of FDA-approved drugs with the potential to alter the pH of endosomes (Figure 4A). We selected a panel of 6 drugs which includes those acting on Na+/K+ ATPase (Omeprazole, Esomeprazole, Pantoprazole, SCH-28080, Lansoprazole) and a protonophore that disrupts proton gradient (Niclosamide). We developed a quantitative high throughput screening pipeline for testing these drugs in both endocytic assay as well as pH estimation assay in AGS cells. The screen was carried out at a concentration of 10µM for all drugs.

**Figure 4:**
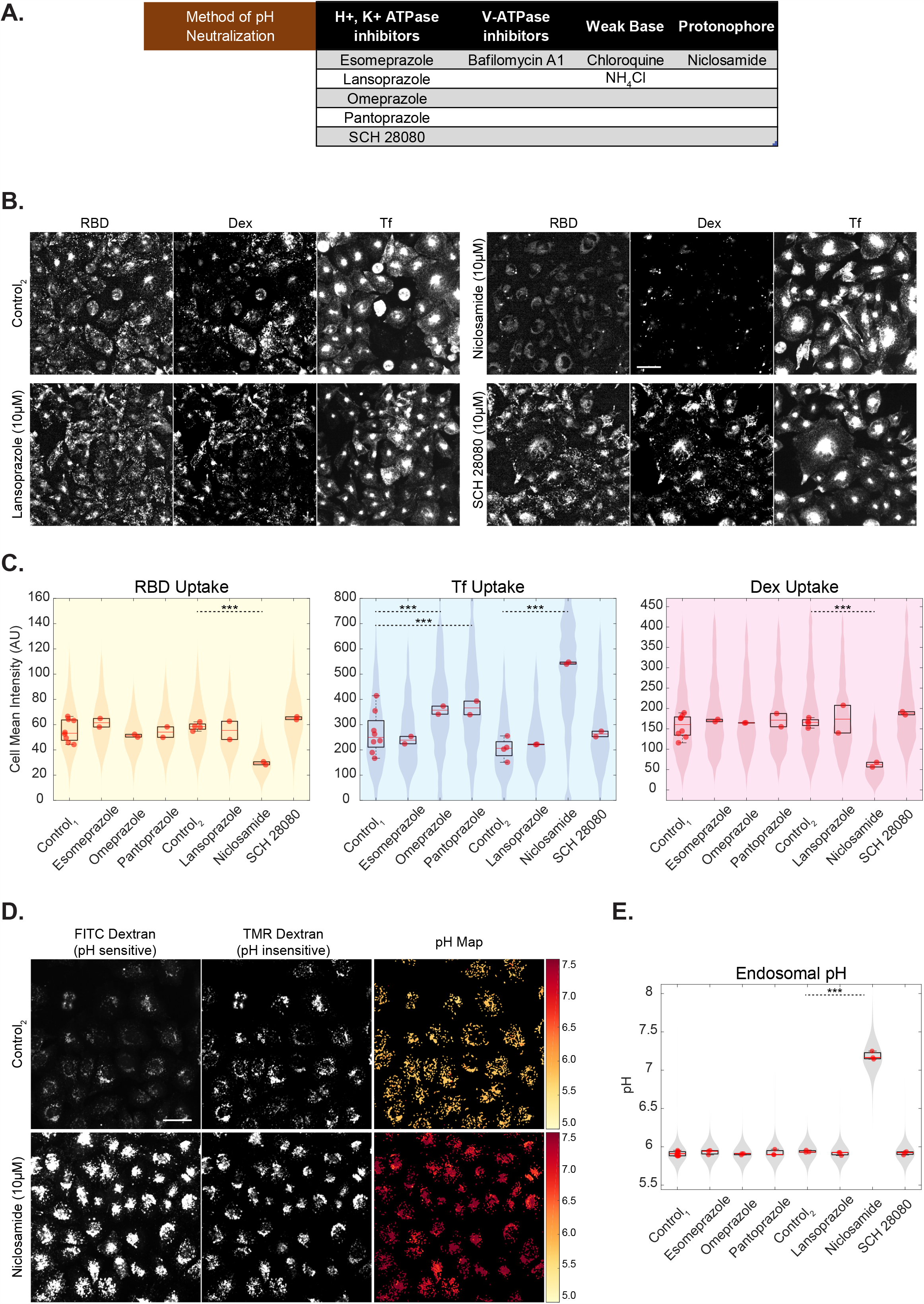
Identifying FDA-approved drugs functioning similar to BafA1 and NH_4_Cl. A: Table describing various methods of altering endosomal pH along with the chosen subset of drugs to screen for entry and acidification inhibition B, C: High-throughput assay in which AGS cells were treated with an array of drugs at 10µM concentration for 30 minutes and pulsed with RBD, dextran and transferrin for 30 minutes. Niclosamide shows reduction in RBD (p-value < e-195) and dextran (p-value < e-133) uptake and increase in transferrin uptake (p-value < e-155). Omeprazole (p-value < e-33) and Pantoprazole (p-value < e-37) show an increase in transferrin uptake and minimally affects RBD or dextran uptake. Images are shown in B (and in S7A), quantification is shown in C and p-value table for all the inhibitor treatments is indicated along with S7A legends. Control_1_ is 0% DMSO and Control_2_ is 0.3% DMSO. Number of repeats ≥ 2 for each treatment and each repeat has >80 cells. D, E: High-throughput assay in which AGS cells were pulsed with FITC and TMR dextran for 2 hours, chased for 1 hour with an array of drugs or control and imaged live. Endosomal pH increases upon addition of Niclosamide (p-value < e-110). Images along with pH maps are depicted in D (and in S7B), quantification is shown in E (and in S7C) and p-value table for all the inhibitors is indicated along with S7C legends. Control_1_ is 0% DMSO and Control_2_ is 0.2% DMSO. Number of repeats ≥ 3 for each treatment and each repeat has >80 cells. Data representation in C and E is as described in Figure 2. Scale bar: 40µm (B, D).

Of the 6 drugs tested in the endocytosis assay, Niclosamide showed the strongest effect on the uptake of the 3 probes (RBD, dextran and transferrin) similar to what we observed for acidification inhibitors. Niclosamide treated cells showed reduced RBD and dextran uptake and increased transferrin uptake (Figure 4B, 4C). It is interesting to note that while the other proton pump inhibitors had minimal effects on RBD or dextran uptake at the concentration tested, Omeprazole and Pantoprazole showed a significant increase in transferrin uptake (Figure 4C, S7A). This suggests that these two drugs could specifically act on the transferrin containing endosomes and not in the compartments of relevance for RBD and dextran uptake, while Niclosamide inhibits the RBD and dextran uptake.

Of the 6 drugs tested in the late endosomal pH estimation assay, Niclosamide also showed the strongest neutralization effect on the pH of acidic endosomes (Figure 4E) by increasing the endosomal ratio of FITC/TMR (Figure S7C). The other drugs had minimal effects on the pH of late endosomes at the concentration tested (Figure 5E, S7B). The spatial pH maps of Niclosamide treated cells show an increase in pH in the majority of endosomes within the cell (Figure 5D). Niclosamide increased the FITC endosomal intensity and reduced the numbers of endosomes (Figure S7C) similar to the effect of BafA1 on these endosomal trafficking parameters.

**Figure 5:**
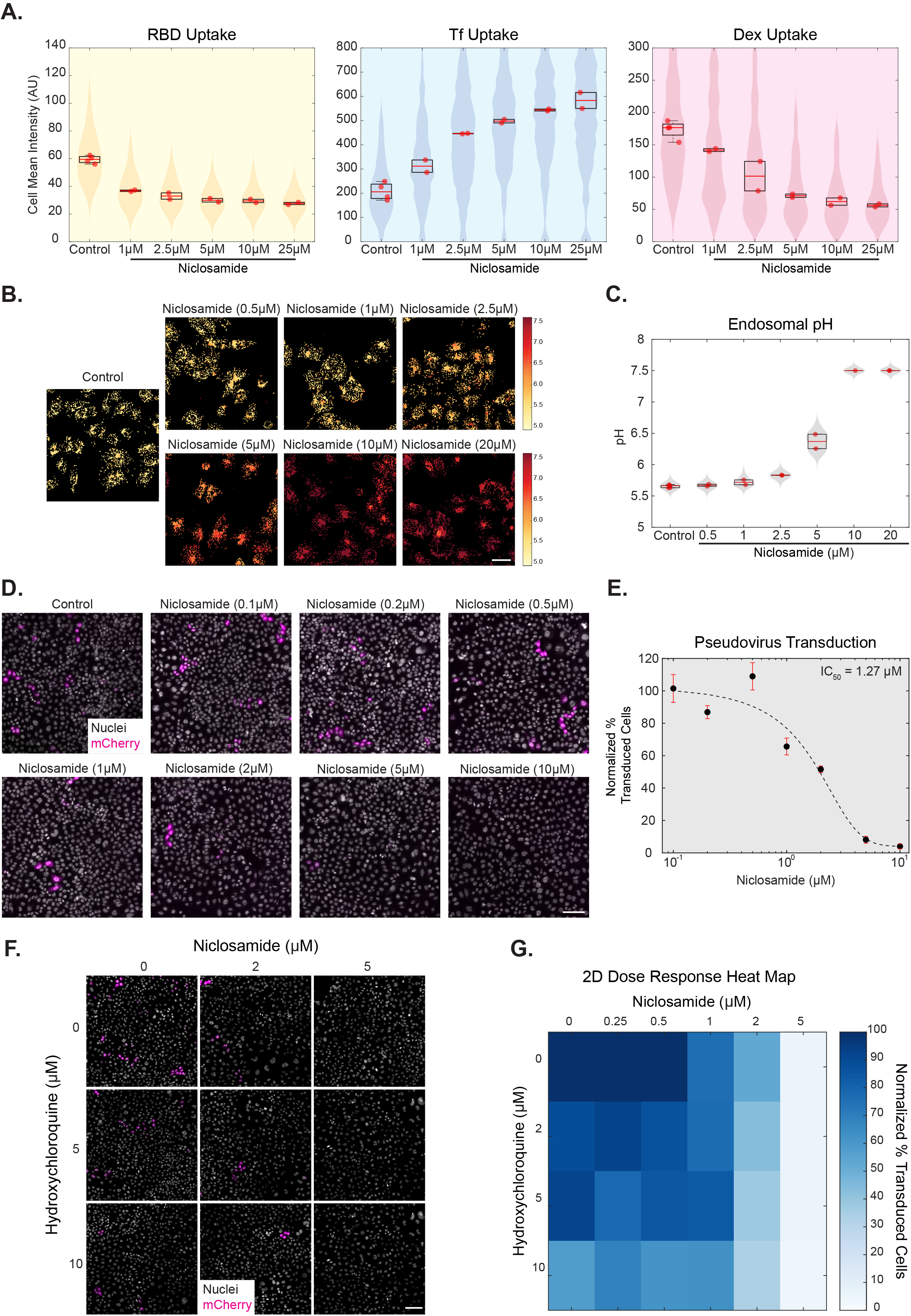
Niclosamide functions as an acidification and entry inhibitor. A: High-throughput endocytic assay in which AGS cells were treated with different concentrations of Niclosamide for 30 minutes followed by pulse of RBD, dextran and transferrin for 30 minutes. RBD and dextran uptake reduces, and transferrin uptake increases in a dose-dependent manner. Images are shown in S8A, quantification in 5A and S8B and p-value table for all the concentrations is indicated along with S8A-S8B legends. Number of repeats = 4 for Control (0.6% DMSO) and 2 each for each concentration of Niclosamide. Each repeat has >80 cells. B, C: High-throughput pH estimation assay in which AGS cells were pulsed with FITC and TMR dextran for 2 hours, chased for 1 hour with different concentrations of Niclosamide and imaged live. A dose-dependent increase in endosomal pH is seen with increasing Niclosamide concentrations. Images along with pH maps are shown in 5B and quantification in 5C and S8C. p-value table is indicated along with S8C legends. Number of repeats = 6 for Control, 2 each for each concentration of Niclosamide and 1 for 10µM Niclosamide. Each repeat has >80 cells. D, E: Spike-pseudovirus transduction assay in which AGS cells were preincubated for an hour with different concentrations of Niclosamide or DMSO and incubated along with virus (MOI = 0.5) for 8 hours followed by the continued presence of 100nM Niclosamide or 0.005% DMSO in the media after removal of the virus until termination. Images of cells expressing the reporter mCherry protein in D and Normalized percentage transduction in E show a dose-dependent reduction in transduction efficiency upon treatment with Niclosamide compared to pooled control from 0.2%, 0.1%, 0.04%, 0.01% and 0.005% DMSO treatments. Black dots with red error bars represent mean +/-SD. Dotted line represents the sigmoidal fit of the means across different Niclosamide concentrations. Refer to S9A(i) for % viability quantification, S9A(ii) for transduction efficiency after 4 hours of incubation with the virus and S9A legends for p-value table. F, G: Images in F and 2-dimensional dose-response heat map in G depict the combinatorial Spike pseudovirus transduction assay in AGS cells with indicated concentrations of Niclosamide (0-5µM) and Hydroxychloroquine (0-10µM). Normalized percentage transduction of cells is represented as a heat map. Refer to S9F for % viability quantification. Data representation in A and C is as described in Figure 2. Scale bar: 40µm (B) and 100µm (D, F).

Omeprazole and other proton pump inhibitors are prodrugs which are used for treating Gastro-esophageal reflux disease (GERD) ^65^. They are activated by low pH, bind covalently to H+/K+ ATPase and inhibit the enzymatic function ^66^. We tested the hypothesis if these drugs could also similarly block the proton pumps in the late endosomes and thus increase the endosomal pH ^67,68^. Earlier studies have indicated that Omeprazole ^69^, Lansoprazole ^70^, and Pantoprazole ^71^, neutralize the endosomal pH only when used at very high concentrations (> 1mM) in EMT-6 and MCF-7 cells. However, the plasma concentration of these proton pump inhibitors varies between 1–23µM ^65^. Thus, at least in the concentration range of relevance, we find no effect of these drugs on the acidification of endosomes and the uptake of RBD.

### Niclosamide functions as an acidification and entry inhibitor

Niclosamide is an anti-helminthic FDA-approved drug and has been in use since the 1960s (Ditzel, 1967). Many recent studies show that Niclosamide has broader clinical applications and has also been identified as an antiviral against SARS-CoV, human Rhinovirus, Influenza viral, Dengue virus ^72,73^. As Niclosamide emerged as a potential drug candidate in both the RBD endocytosis as well as endosomal pH neutralization screens, we investigated the dose-dependent role of Niclosamide in reducing RBD uptake, neutralizing endosomal pH and inhibiting Spike-pseudovirus infection. We found that Niclosamide reduced both RBD and dextran uptake, as well as increase transferrin uptake in a dose-dependent manner (1–25µM) (Figure 5A, S8A). We observed Niclosamide’s effect on RBD endocytosis even at concentrations as low as 1µM. On analyzing the effect of Niclosamide on endosomal numbers and intensity, we found that Niclosamide increased the endosomal intensity of transferrin endosomes and reduced the number of RBD and dextran endosomes (Figure S8B). These effects are remarkably similar to the effects observed with acidification inhibitors – BafA1 and NH_4_Cl. We also confirmed the inhibitory effect of Niclosamide on RBD and dextran uptake in another cell line – HEK-293T (Figure S10A-D), and on normalized RBD uptake in AGS cells overexpressing ACE2 (Figure S3C, S3D).

Further, we also observed a dose-dependent effect of Niclosamide on neutralizing the pH of late endosomes, with neutralization effects seen even at 2.5µM (Figure 5B, 5C). The dose-response effect is seen on the ratio of endosomal FITC/TMR as well as other endosomal trafficking parameters - FITC and TMR endosomal intensities and numbers of endosomes (Figure S8C). The spatial pH maps of cells also show a gradual shift of endosomal pH from acidic to neutral pH with different doses of Niclosamide (Figure 5B), especially at 2.5µM wherein some endosomes within the cell are still acidic while some others are neutralized. Towards evaluating the effect of Niclosamide on the pH of early time point endosomes, AGS cells were labelled with FITC and TMR dextran for 20 minutes and chased for 10 minutes with or without Niclosamide for the entire duration of pulse and chase (Figure S4E). Unlike BafA1, while Niclosamide reduced the net uptake of dextran, similar to BafA1, Niclosamide also increased the endosomal FITC intensity and endosomal FITC/TMR ratio of early time point (30 minutes) endosomes (Figure S4F), indicating that Niclosamide neutralizes the pH of these endosomes as well.

We assessed the effect of different concentrations (0.1-10 µM) of Niclosamide on Spike-pseudovirus entry in AGS cells, using the experimental strategy designed to assess virus entry as described before. At the end of 4 and 8 hours of viral incubation, the pseudovirus containing media along with the Niclosamide was removed and all treatments were replenished with media containing a reduced concentration of 0.1µM Niclosamide. We observed a strong dose-dependent reduction of transduction efficiency as a function of increasing Niclosamide concentration at both viral incubation durations with negligible toxicity (Figure 5D, S9Ai and Aii). IC_50_ of ∼1.27µM was estimated on fitting a sigmoidal function to the dataset obtained for 8 hours of viral incubation (Figure 5E, Methods). Together, all three assays conclude that Niclosamide can act as acidification and entry inhibitor.

### Enhancing the inhibition of infectivity: A combination strategy with Niclosamide and Hydroxychloroquine

A combinatorial approach of drugs with varying mechanisms of inhibition works as an effective therapy to combat infection ^74^. Given that Niclosamide exhibits a short half-life ^75^, has poor bio-availability (∼10 %) ^76^ and our observations indicate moderate IC_50_ for inhibition of Spike-pseudovirus transduction, we tested if the action of Niclosamide can be enhanced in the presence of another FDA-approved drug known to be effective against SARS-CoV2 infection. Since published reports and commonly practiced treatments against SARS-CoV2 infection employ Hydroxychloroquine (HCQ), a less toxic variant of Chloroquine ^77^, we tested the effect of HCQ on altering late endosomal pH and Spike-pseudovirus transduction assay. Like Chloroquine, cells treated with 50µM HCQ also minimally altered the late endosomal pH (Figure S9B, S9C). However, we observed the pseudovirus transduction to be markedly reduced at HCQ concentrations of 50µM and 25µM (Figure S9D, S9E) and only modestly reduced at the concentrations of 10µM or lower in a dose-dependent manner (See the first box plot in Figures S9 Fi, Fii and Fiii). To assess the synergistic effect of the two drugs, we chose a concentration range with the maximum concentrations of 10µM HCQ and 5µM Niclosamide. The 2-dimensional dose-response map shown in Figure 5G summarizes the effect of the two drugs on transduction. We observed an augmented reduction in infection when HCQ was used at a concentration of 10µM along with varying concentrations of Niclosamide compared to where HCQ was used at 0, 2 and 5µM (Figure S9Fi, Fii and Fiii). These results indicate an additive effect on inhibition of pseudovirus transduction when effective concentrations of HCQ is added along with effective concentrations of Niclosamide (Figure 5F, 5G). Thus, Niclosamide could potentially enhance the efficacy of the plethora of treatments currently being used to combat SARS-CoV2 infection.

## Discussion

Understanding the molecular mechanisms of viral entry into target cells is critical to design effective treatments and prevention strategies against infection. Employing various methodologies, we report for the first time that fluorescently labelled RBD of SARS-CoV2 enters cells through a pH-dependent CG pathway. High-resolution quantitative imaging approaches enabled us to detect the localization of RBD to acidic compartments. Endosomal acidification inhibitors that affect the uptake of CG cargo also inhibit RBD uptake. Complementing our observations with RBD, we show that infection by Spike-pseudovirus is also dependent on endosomal acidification. Further, by employing a targeted drug screen, we have identified Niclosamide as a potential inhibitor against SARS-CoV2 entry (Figure 6).

**Figure 6:**
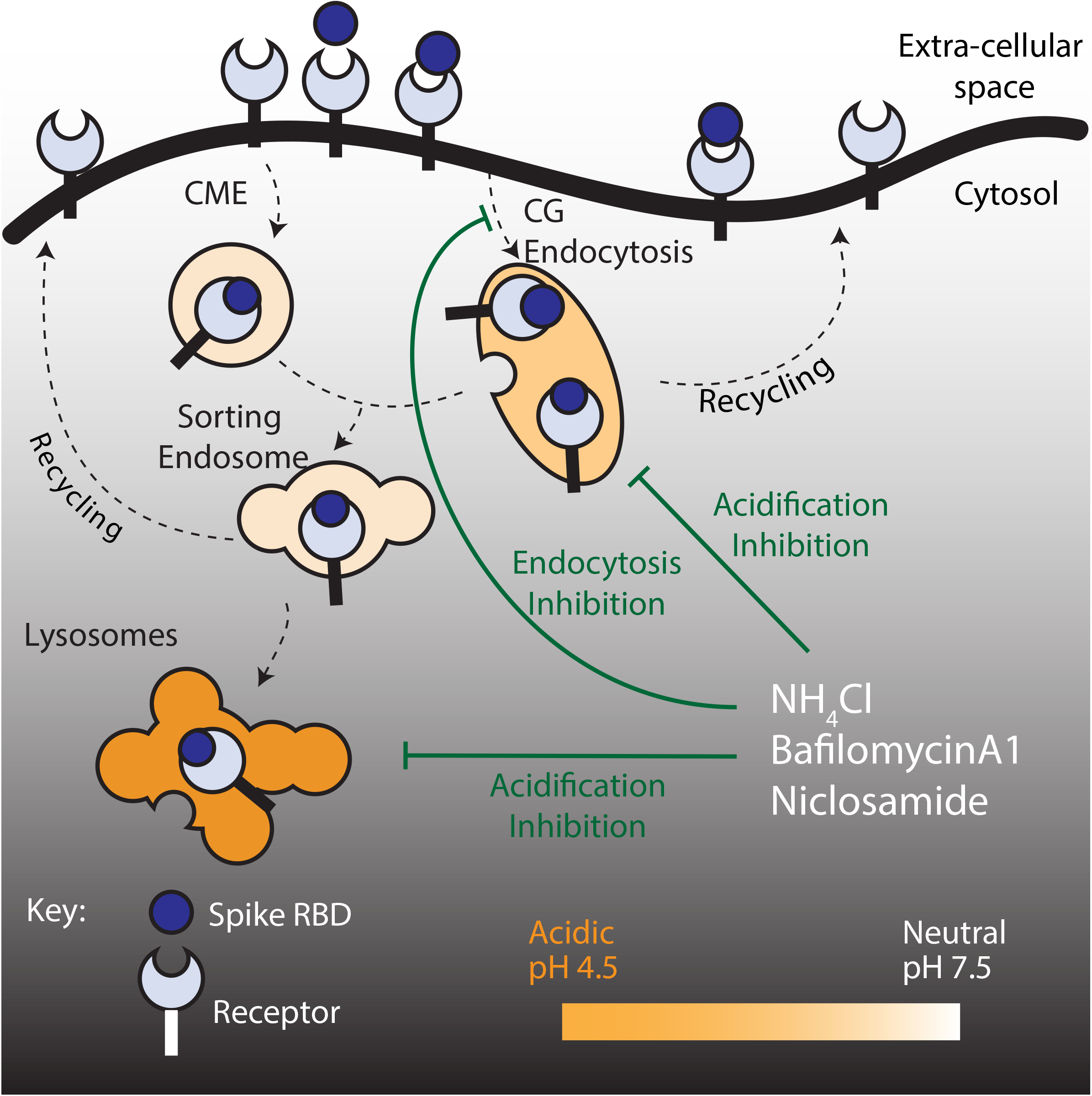
Model. Spike RBD interacts with receptors at the cell surface and is internalized via the CLIC/GEEC pathway. Acidification inhibitors neutralize the pH of endosomes as well as block entry via the CG pathway. Niclosamide, a protonophore and an FDA-approved drug, also increases the pH of endosomes and blocks uptake via the CG pathway. Spike pseudovirus infection assays confirm that acidification inhibitors, including Niclosamide, prevent viral transduction.

The choice of viral entry into host cells is influenced by cell surface interacting partners and co-factors ^11,25^. Although ACE2 has been identified as the receptor for SARS-CoV2, other receptors are being uncovered. These include Neuropilin ^12,13^, CD147 ^14^, Heparan Sulphate proteoglycans ^15^ and HDL scavenger receptors ^16^. Additionally, the highly glycosylated nature of Spike protein could also confer the ability to interact with yet unidentified receptors. These virus-receptor interactions could potentially dictate the endocytic route employed by the virus. This is exemplified by our observation that although RBD uptake is reduced upon blocking the CG pathway, residual RBD re-routes towards the CME and enables pseudovirus infection. Re-routing could presumably be due to binding to different receptors that could follow alternative internalization routes. Whether the Spike-pseudovirus follows routes of entry like RBD, can be addressed with tractable pseudoviruses or synthetic virus-like particles. However, recent genome wide screens ^38,78^ indicating the importance of cholesterol homeostasis in SARS-CoV2 infection are consistent with a cholesterol senstitive CG endocytic route ^79^ in entry. Knockouts of genes affecting cholesterol biosynthesis (SCAP, MBTSP1, MBTSP2) not only reduced infection of native SARS-CoV2 but also of Spike-pseuodviruses indicating that cellular cholesterol is necessary for efficient Spike mediated entry of SARS-CoV2 ^78^.

Known inhibitors of endosomal acidification, BafilomycinA1 and NH_4_Cl, play an important role in neutralizing acidic lysosomes and thus subverting viral membrane fusion and entry of several viruses ^11,24–27^. Here, we report that these inhibitors also play a more upstream role by inhibiting the endocytosis of RBD itself. Both these treatments inhibited the uptake of CG cargo and RBD, reduced Spike-pseudovirus infection and drastically elevated endosomal pH. It is interesting to note that the inhibition of acidification in addition to dramatically reducing CG uptake did not cause re-trafficking of RBD through another endocytic pathway, as was observed for other CG inhibitors. This suggests that the acidification inhibitors could negatively influence the RBD-receptor interactions at the cell surface along with further ramifications of blocking the CG pathway.

These observations encouraged us to screen a subset of FDA-approved compounds known to affect endosomal acidification: proton-pump inhibitors (Omeprazole, Lansoprazole, Pantoprazole, Esomeprazole, SCH-28080), and protonophore (Niclosamide). Of all the 6 compounds tested only Niclosamide inhibited CG cargo and RBD uptake, elevated endosomal pH and concomitantly inhibited Spike-pseudovirus infection, all in a dose-dependent manner with an IC_50_ of 1.27 µM in AGS cells. Among several mechanisms of action ^75^, Niclosamide disrupts proton gradient across mitochondrial ^80^ and endosomal ^72^ membranes. The elevated endosomal pH brought about by Niclosamide was shown to inhibit human rhinovirus infection ^72^. Additionally, Niclosamide has been identified as an anti-viral agent against SARS ^81^, Dengue ^73^, MERS ^82^ and more recently proposed for SARS-CoV2 (with IC_50_ of 0.28 µM in Vero cells) ^83^, although the mechanism of action remained unknown. In contrast, the proton pump inhibitors used in our study failed to interfere with RBD uptake. This could be because they remained inactive ^65^ or the concentrations tested predominantly affect H+/K+ ATPases, while mM concentrations are required to inhibit V-ATPases ^67^. Along these lines, studies show that proton pump inhibitors inhibit Ebola-pseudovirus ^84^, SARS-CoV and SARS-CoV2 ^85^ infection only when used beyond achievable plasma concentrations ^65^.

Surprisingly, Chloroquine did not affect RBD uptake and only marginally raised the endosomal pH. However, it caused a strong inhibition of Spike-pseudovirus infection. This strongly suggests that Chloroquine could be operating in the initial steps of viral infection but post endocytosis ^64^, as observed with RBD uptake. Chloroquine is likely to function in many pH-independent ways to inhibit SARS-CoV2 infections, distinct from Niclosamide. For example, by altering terminal glycosylation of ACE2 ^86^; via its activity as a zinc ionophore affecting ACE2 activation ^87,88^; by interacting with ER resident Sigma receptors that initiates cell stress response ^89^; by its ability to strongly bind a viral protease essential for Spike activation ^90^. At this time, the exact mechanism(s) by which Chloroquine inhibits SARS-CoV2 entry remains unclear.

In conclusion, our study reports the high capacity CG pathway as a potential endocytic route for SARS-CoV2. We further show that endosomal acidification is critical for SARS-CoV2 entry and infection and can be a promising therapeutic target as observed by the results seen with Niclosamide, BafilomycinA1 and NH_4_Cl. This study also paves way for large-scale screens to repurpose FDA-approved drugs as acidification inhibitors and scrutinize for more Niclosamide-like drugs that might have better bioavailability or can be used in combination with other antiviral drugs. Moreover, the methods described in our study can effectively be extended and better represented with clinical isolates of viruses to assess their infective journey in primary cells that represent the more natural hosts for infection.

## Materials and Methods

### Cell lines, constructs, and antibodies: See supplementary methods for more details *Chemicals and reagents*

Niclosamide and AN96 were chemically synthesized and proton pump inhibitors, Esomeprazole and Pantoprazole, were extracted from commercially available tablets as detailed in the Supplementary Methods. The other proton pump inhibitors, Lansoprazole and SCH-28080, were obtained from the LOPAC®1280 library, and Omeprazole was procured from Sigma (O104).

### Endocytosis assays

AGS or HEK-293T cells were plated in 35mm coverslip bottom dishes and processed after 48 hours at 60-70% confluency. Cells were washed twice with HEPES buffer (wash and imaging buffer composition: 150mM NaCl, 20mM HEPES, 5mM KCl, 1mM CaCl_2_, 1mM MgCl_2_, 2mg/ml Glucose, pH 7.5) at 37°C. Endocytosis was monitored using fluorescently labelled RBD (Alexa/Atto 488, 10μg/ml), 10kDa TMR-dextran (1mg/ml) and/or Iron-loaded Transferrin (10μg/ml, Alexa 647) in serum-free medium for indicated time points at 37°C. Endocytosis was stopped using ice-cold wash buffer and cells were subsequently fixed with 2.5% paraformaldehyde (PFA) for 20 minutes at room temperature (RT). Cells were then washed and imaged. For inhibitor experiments, cells were pre-treated with various inhibitors (AN96 25μM, ML141 50μM, Amiloride 1mM, BafA1 200nM or 400nM, NH_4_Cl 30mM) and respective controls in serum-free medium for 30 minutes at 37°C and inhibitors were maintained during endocytic assays.

To measure normalized transferrin or normalized RBD uptake (Figures 1H-1I, S3C-S3D), cell surface-bound probes after the endocytic pulse with transferrin or RBD were stripped using two washes with ice-cold ascorbate buffer (160mM sodium ascorbate, 40mM ascorbic acid, 1mM MgCl_2_, 1mM CaCl_2_, pH 4.5), followed by three washes with ice-cold wash buffer at 4 °C. Cells were then fixed with ice-cold 2.5% PFA for 5 mins at 4 °C and 15 minutes at RT. Transferrin receptor (TfR) was labelled by incubating cells with fluorescently labelled anti-hTfR (OKT-9) for 2 hours at RT. To label surface ACE2, fixed cells were blocked with 10mg/ml bovine serum albumin (30 minutes) followed by incubation with anti-myc primary antibody (1 hour) and secondary antibody (45 minutes) in blocking buffer at RT. Cells were then washed and imaged.

### pH estimation assays

For estimating the pH of late endosomes, cells were pulsed with pH-sensitive 10kDa FITC-dextran (1mg/ml) and pH-insensitive 10kDa TMR-dextran (1mg/ml) for 2 hours in serum-free media, chased for 1 hour in the presence of inhibitors or control and imaged live. The above pulse and chase times were chosen to allow the accumulation of labelled dextran in acidic late endosomal and lysosomal compartments (co-labelled with Lysotracker, data not shown). To estimate the endosomal pH, the ratio of FITC to TMR fluorescence was computed and compared to a pH calibration curve (Figures S4A-S4B) which was generated by equalizing the endosomal pH to that of an external buffer. After the pulse with FITC and TMR-dextran and chase, cells were incubated with 5µg/ml nigericin containing buffers of different pH for 10 minutes and imaged to evaluate FITC/TMR ratios for each pH.

For estimating the pH of late endosomes using the 488/458 excitation ratio of FITC-dextran (Figures S5E-S5F), cells were pulsed with FITC-dextran at 1mg/ml for 2 hours, followed by chase in the presence or absence of inhibitors and imaged live.

For estimating the FITC/TMR ratio of early endosomes (Figures S4E-S4F), cells were incubated with pH-sensitive 10kDa FITC-dextran (1mg/ml) and pH-insensitive 10kDa TMR-dextran (1mg/ml) for 20 minutes, chased for 10 minutes and imaged live. Throughout the pulse and chase duration, the cells were incubated in serum-free media with control (0.2%DMSO) or BafA1 400nM or Niclosamide 10µM.

### Spike-pseudovirus transduction assays

AGS/HEK-293T cells were plated in optical bottom 96-well plates. 36 hours post-plating, when cell numbers were ∼4000, transduction was carried out at indicated MOIs. For inhibitor treatment, cells were pre-incubated with indicated concentrations of NH_4_Cl/ BafA1/ CQ/ Niclosamide/ HCQ/ AN96/ ML141, for 1 hour. This was followed by addition of the Spike-pseudoviruses in presence or absence of the inhibitors. At the end of 2/4/8hours, media containing pseudoviruses and inhibitors was removed, and cells were washed once with drug-free media. This was followed by addition of media with or without inhibitor: NH_4_Cl, BafA1 and CQ were removed from the media; Niclosamide, AN96 and HCQ were maintained at a low concentration of 100nM, 1µM and 500nM respectively. This was done to assess the effects of the inhibitors at the initial stages of inhibition, minimize long-term toxicity to the cells as well as to avoid effects on the translational processes of the reporter gene post entry. After 60 hours, cells were fixed, nuclei were labelled with Hoescht and assessed for transduction efficiency based on mCherry reporter expression. In the case of HEK-293T cells (Figure S10G), MTT cell viability assay was performed to check toxicity (assay described in Supplementary Methods).

### Imaging and Analysis

#### a. Endocytic and pH estimation assays

For 35mm dish-based endocytic experiments, fixed samples were imaged using confocal microscopy (Olympus FV3000, 20X/0.85NA objective) to image RBD, dextran and transferrin endosomes with Z sections of 1µm. Maximum intensity projected images were used for further analysis. Cell ROIs were drawn and features such as cell mean intensity in each channel was extracted. For high-throughput endocytic and pH estimation experiments, automated imaging (Spinning disc, Phenix Perkin Elmer, 40XW/1.1NA objective) was used to image nucleus along with RBD, dextran and transferrin (for endocytosis) or FITC and TMR dextran (for pH) with Z sections of 1µm each. For both assays, cell profiler based pipeline was used to segment cells, nucleus and endosomes and extract features as described in supplementary methods. For pH calibration, the mean of the endosomal ratio distributions at different extracellular pH was fit to a sigmoidal equation. For both assays, custom MATLAB routines were used to estimate the endosomal intensities, the number of endosomes and cell mean intensities. In addition, for pH assays, endosomal ratio (FITC/TMR) and endosomal pH (using the calibration curve) for each endosome was computed. As the endosomal intensity distribution within cells is a heavy right-tailed distribution, median endosomal intensity for each probe for each cell was estimated. The distributions of cell mean intensity/endosomal intensities/numbers of endosomes per cell per treatment (for endocytosis) and endosomal intensities/ratio/pH per cell per treatment (for pH) is represented in each quantification.

For 488/458 endosomal ratio estimation experiments, live imaging was done using confocal microscopy (Zeiss LSM 780, 40X/1.4NA objective). Excitation lasers 488nm, 458nm were used and emission was detected using a spectral detector (490nm-560nm). Images were processed as described above to estimate endosomal intensities and endosomal ratios per cell.

#### b. Colocalization analysis

Confocal microscopy (Olympus FV3000, 60X/1.42NA objective) with Z sections of 0.4µm each was employed to image cells across all channels. A MATLAB routine was written to extract colocalization indices. For each cell, endosomes in each channel were segmented based on threshold values. The segmentation in each channel was made finer using morphological operations (dilation followed by erosion). Segmented endosomes were considered for colocalization analysis. Manders’ coefficients and Pearson’s correlation coefficients were computed as described before ^91^.

#### c. Pseudovirus transduction assays

Automated imaging (Widefield, Phenix, 10X/0.3NA objective) of 96 well assay plates was used to image nucleus as well as mCherry positive cells. A cell profiler based pipeline was used to segment nucleus and extract features, as described in supplementary methods. Approximately 50,000 nuclei (cells) were scored for each treatment. A MATLAB routine was written to estimate the % transduction. Mean intensities of the segmented nucleus in the nuclei channel and the mCherry channel for each nucleus across all fields were extracted. Each assay plate included “No-Virus” negative control. This control was used to estimate the background intensities of the mCherry channel within each segmented nucleus. The median of this distribution was considered the background. All nuclei with mCherry intensities of at least 1.8 - 2.2 times (empirically determined) the intensity of the background were considered positive. For each field, the fraction of positive nuclei to the total number of nuclei was determined. The mean of % transduction across all fields for each treatment was calculated. The % transduction was normalized to that of the control and is represented in all the quantifications. The total number of nuclei for each treatment is also represented to understand the effect of the toxicity of drugs.

### Statistical methods and hypothesis testing

All statistical tests between control and treatment were performed in MATLAB using Wilcoxon rank-sum test and the p-value of the hypothesis testing and the number of repeats is indicated in figure legends. In the entire manuscript, ***, **, * and ns indicate p-value of Wilcoxon rank-sum test < 0.001, <0.01, <0.05 and not significant, respectively. It is to be noted that p-value is affected by both the magnitude of differences as well as the sample size. For large sample size, as in case of our high throughput experiments, the impact of random error in measurement will be reduced and the larger magnitude of difference between the control and treatment will be associated with a much smaller p-value.

## Acknowledgements

We thank Nevan Krogan (UCSF, USA) for Spike expression construct, Florian Krammer (Mt. Sinai, USA) for secreted RBD expression construct, Biocon, Ltd (India) and Raghavan Varadharajan (IISc, India) for providing purified RBD for initial experiments, Minhaj Sirajuddin (inSTEM, India) for mCherry expression plasmid, Vinoth Kumar (inSTEM, India) for discussions on Spike-pseudovirus characterization and Mylan Laboratories (India) for providing HCQ. We thank the Central Imaging and Flow Cytometry Facility (CIFF) and the Screening Facility at NCBS, for imaging and high content screening. We sincerely thank Abrar Bhat (for help with RBD purification), Chandrima Patra (for help with imaging), Greeshma Pradeep (for help with microscopes), Sarayu Beri (for help with cell culture) and all members of SM lab for exciting discussions on this project. We acknowledge the technical, administrative and hospitality staff at NCBS for tremendous help especially during the national lockdown. We acknowledge NCBS-TIFR graduate fellowship (for CP and PS), UGC graduate fellowship (for RG) and NCBS postdoctoral fellowship (for SJ). TSvZ acknowledges EMBO postdoctoral fellowship (ALTF 1519-2013) and NCBS Campus fellowship; AC thanks India Alliance DBT – Wellcome Trust Early career fellowship (IA/E/15/1/502339); SM acknowledges J.C. Bose Fellowship from DST, Government of India, and India Alliance DBT – Wellcome Trust Margdarshi fellowship (IA/M/15/1/502018). CP, PS and SJ received support from Margadarshi fellowship grant (IA/M/15/1/502018).

## Author Contributions

CP, RG, PS, AG, VS and SM conceived the study. CP, RG, PS, SJ, AG, VS and SM designed the experiments. Endocytosis and pH assays were done by RG and CP, Spike-pseudovirus assays were done by PS and SJ. CP, TSvZ and DS acquired and analyzed data. CP, RG, PS, SJ, TSvZ, DS, AG, VS and SM interpreted the data. RG, NS, SML, DS and CP conducted high-throughput experiments. AC and PS prepared Spike-pseudovirus. VKN and SBB synthesized Niclosamide. RA, AHN, PPS and RV synthesized AN96. TPP and PV extracted Esomeprazole and Pantoprazole. CP, SD, BM and PS purified and labelled RBD. CP, RG, PS, SJ, TSvZ and SM wrote the manuscript with comments from AC, AG and VS.

