## Supplementary figures and images for "Niclosamide inhibits SARS-CoV2 entry by blocking internalization through pH-dependent CLIC/GEEC endocytic pathway"

### Figure S1

#### Supplementary Figure 1

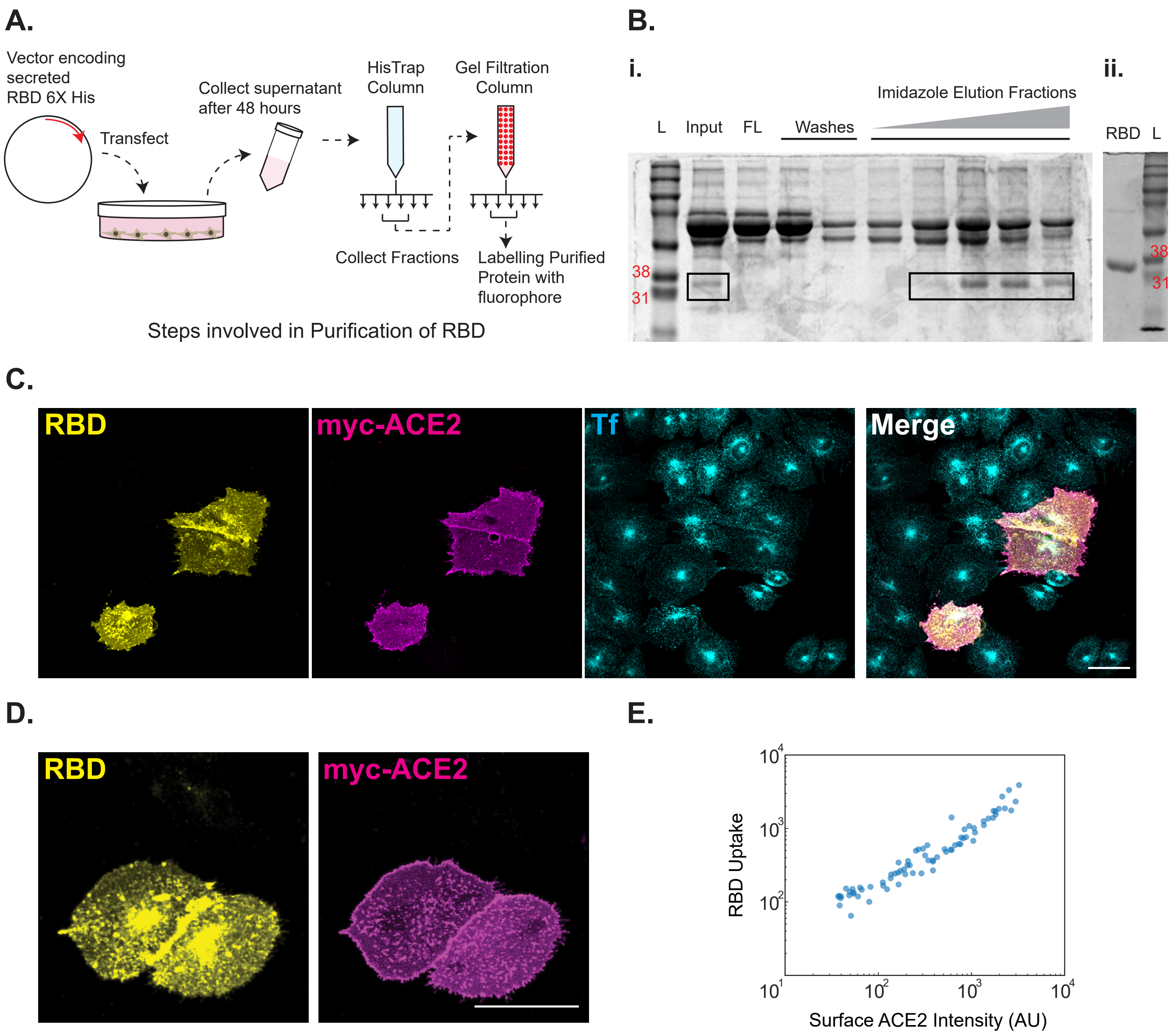

### Figure S2

##### Supplementary Figure 2

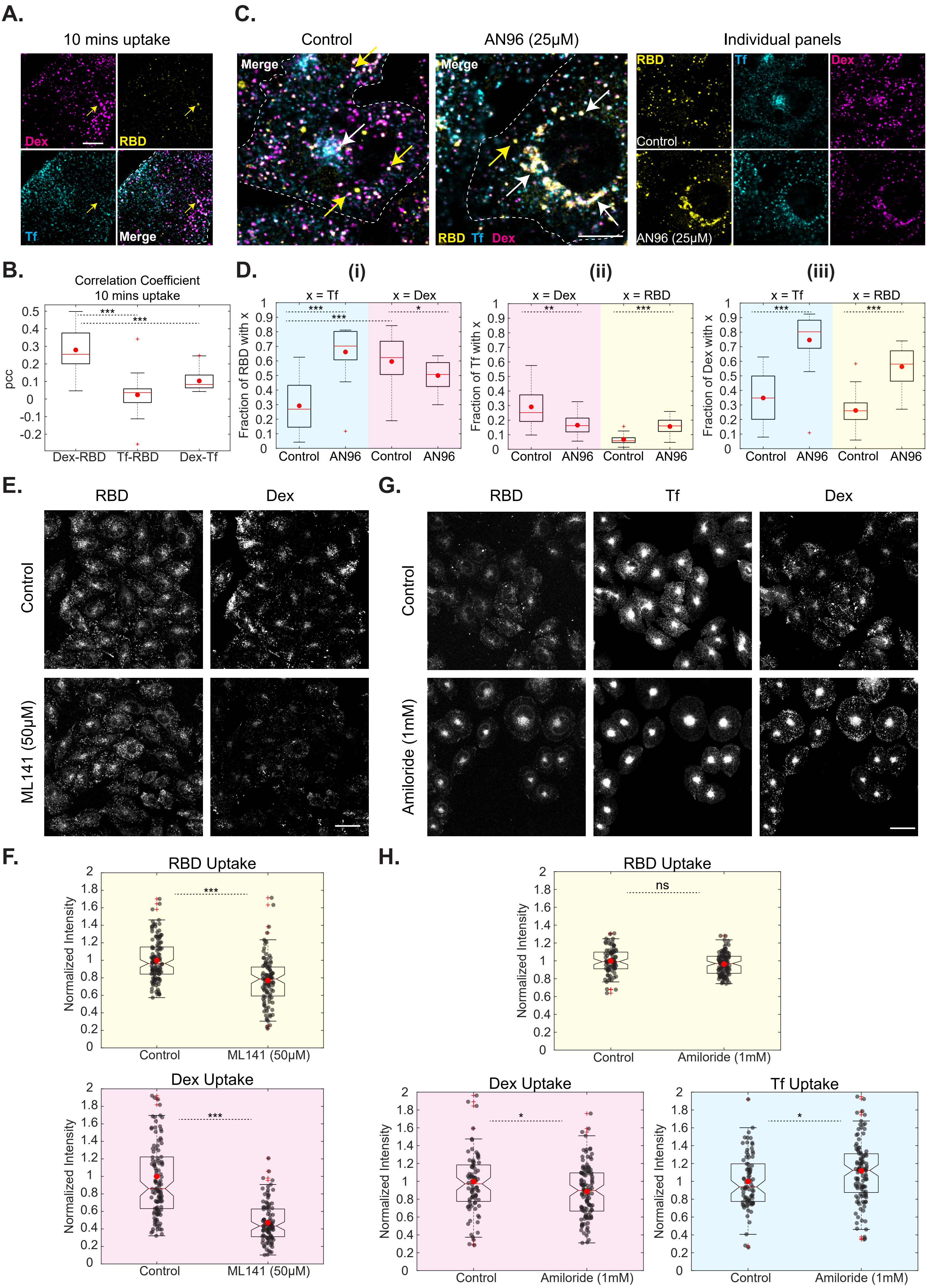

### Figure S3

#### Supplementary Figure 3

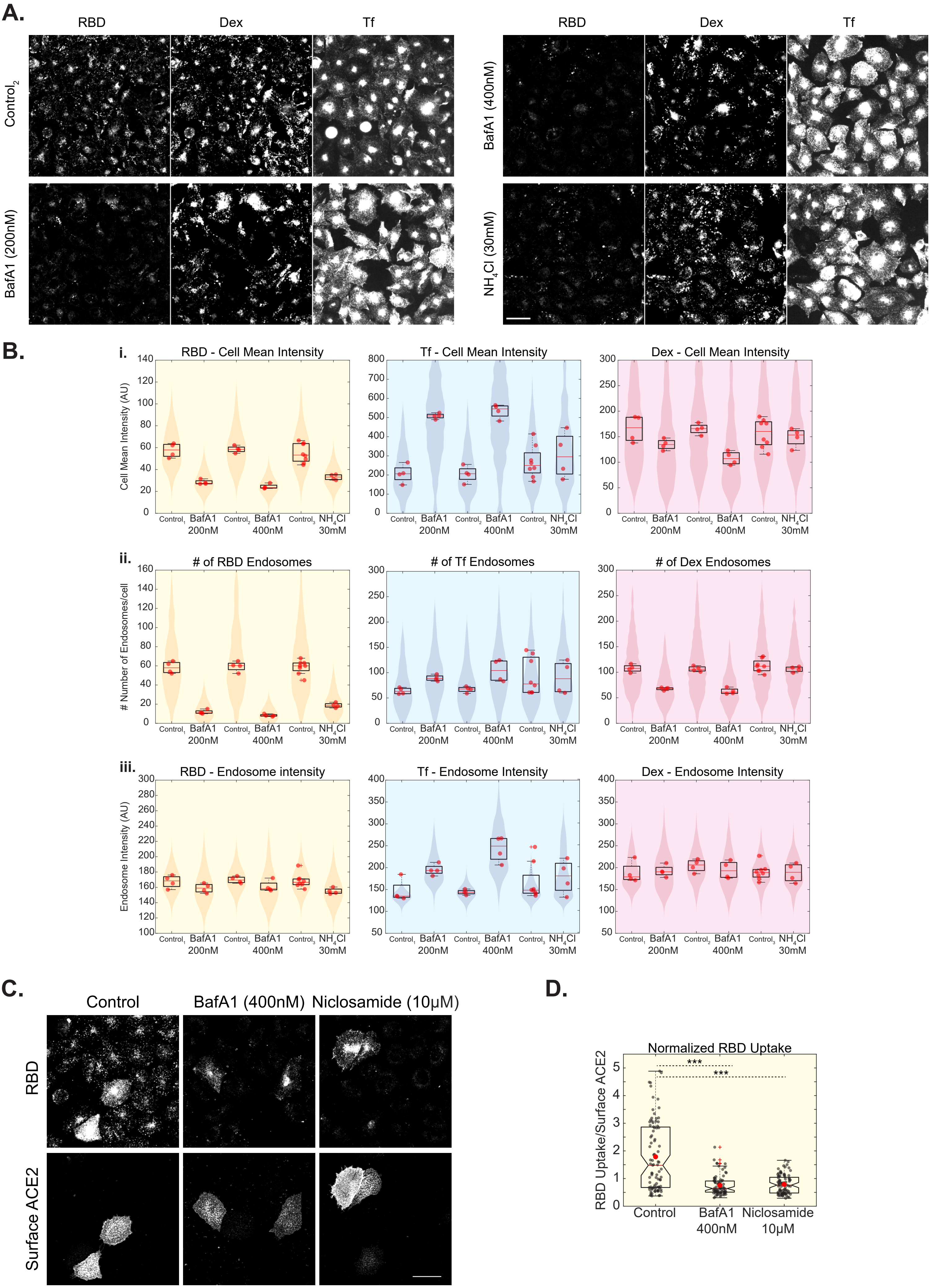

### Figure S4

#### Supplementary Figure 4

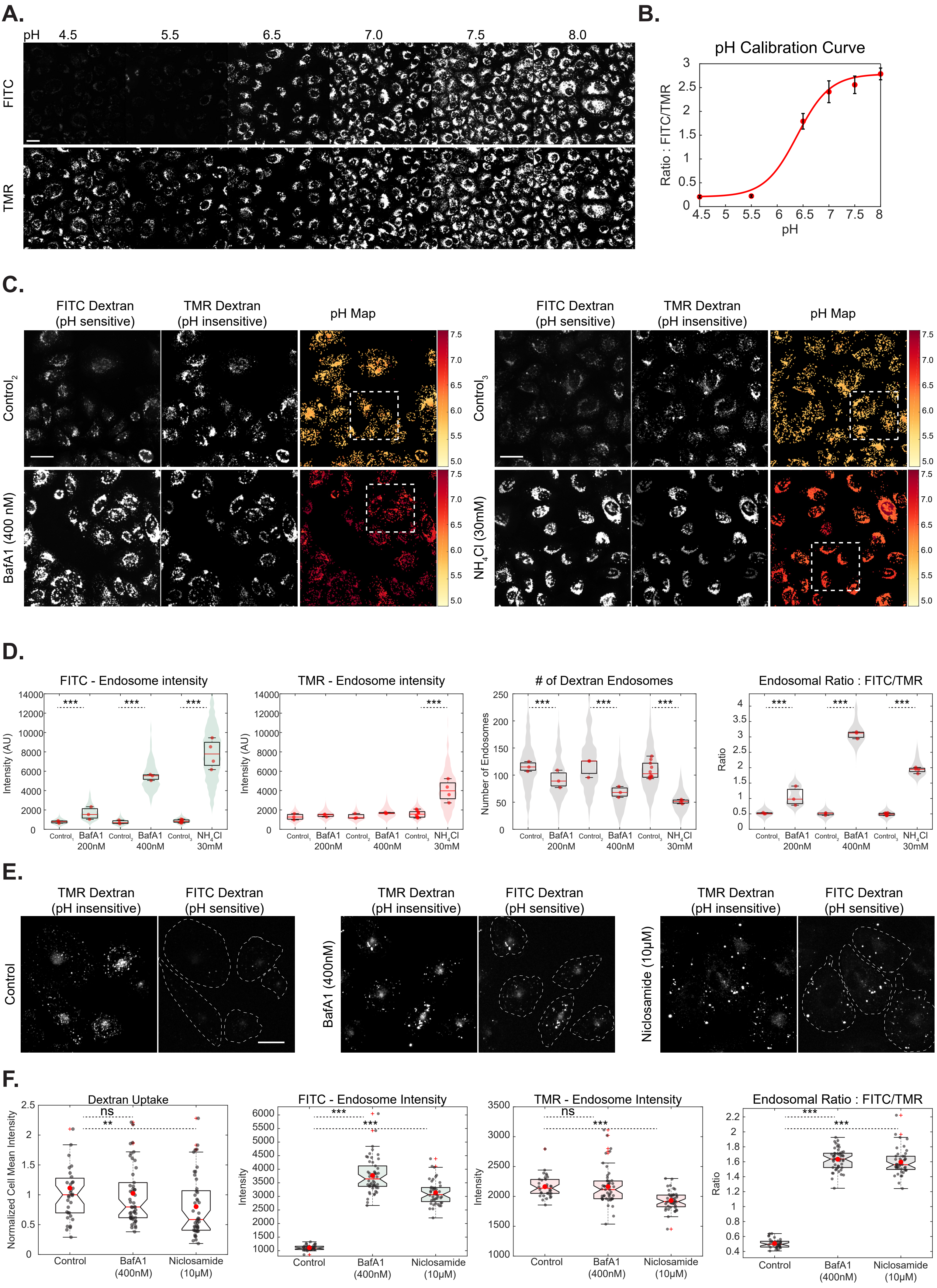

### Figure S5

#### Supplementary Figure 5

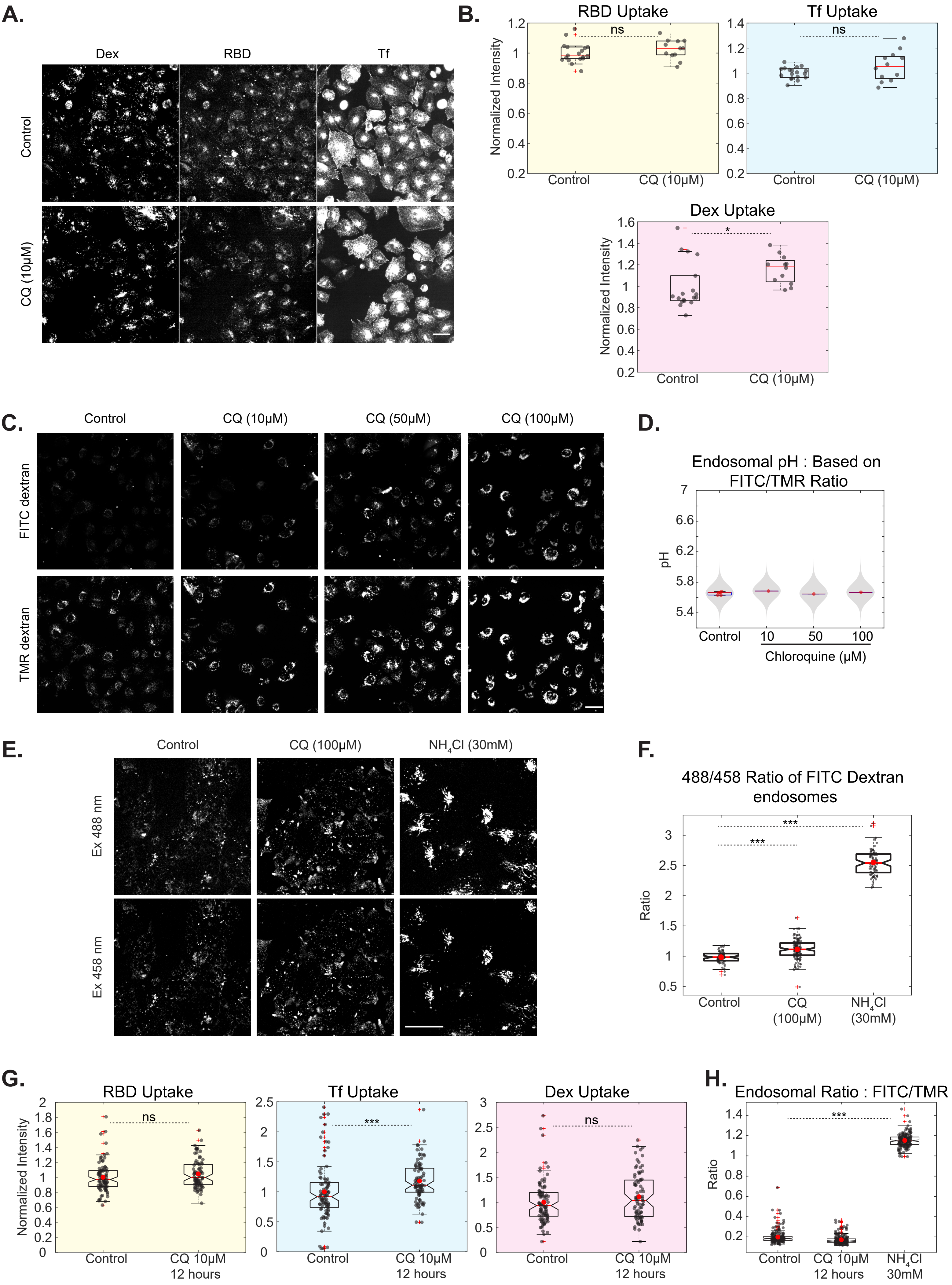

### Figure S6

#### Supplementary Figure 6

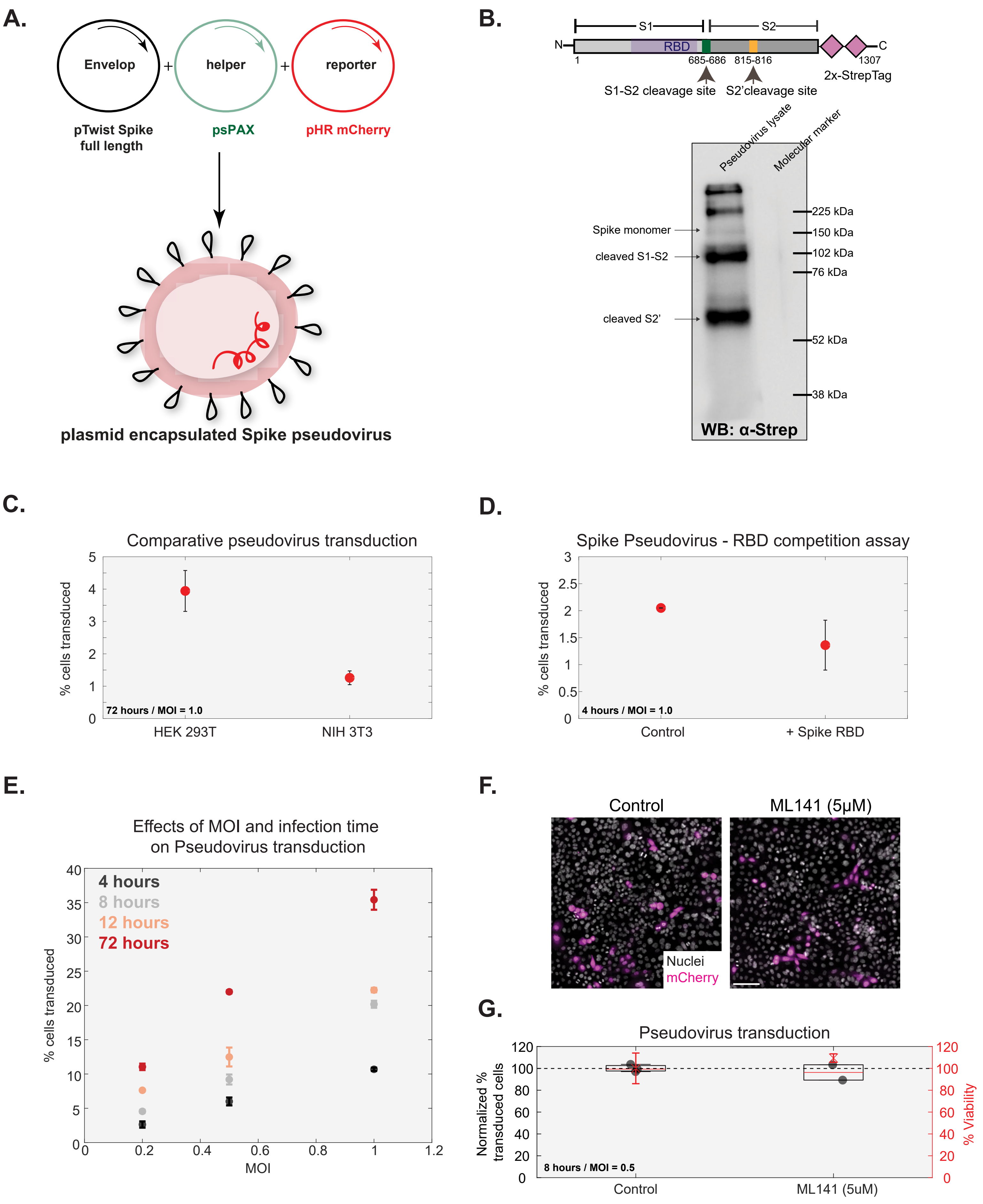

### Figure S7

#### Supplementary Figure 7

A.

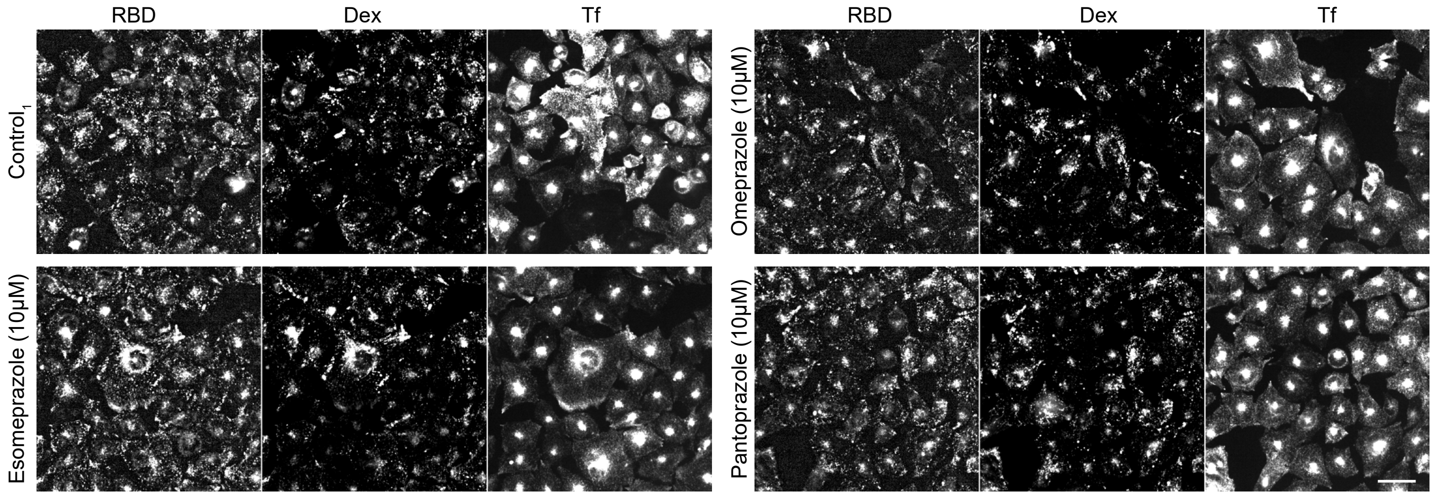


B.

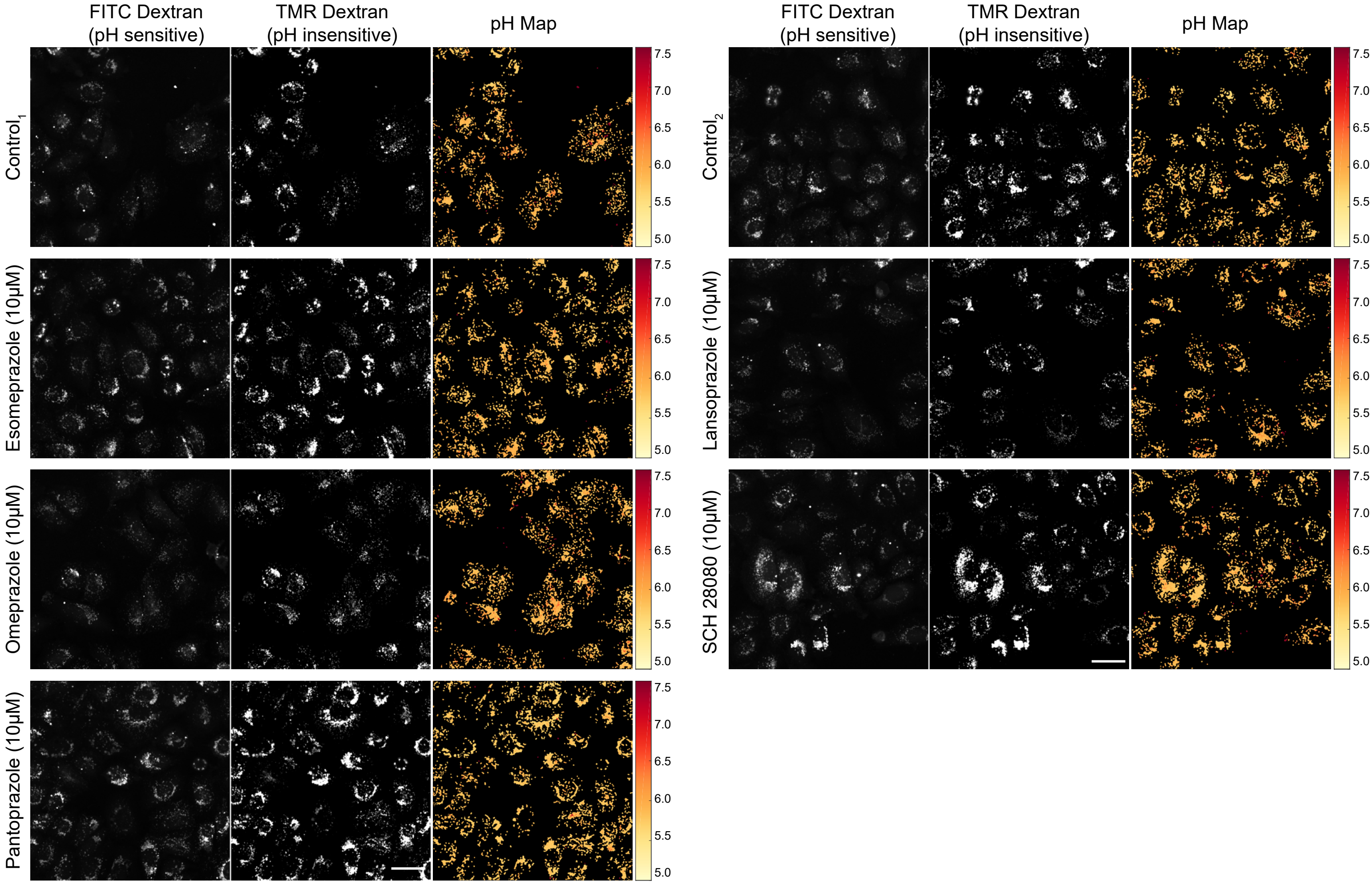


C.

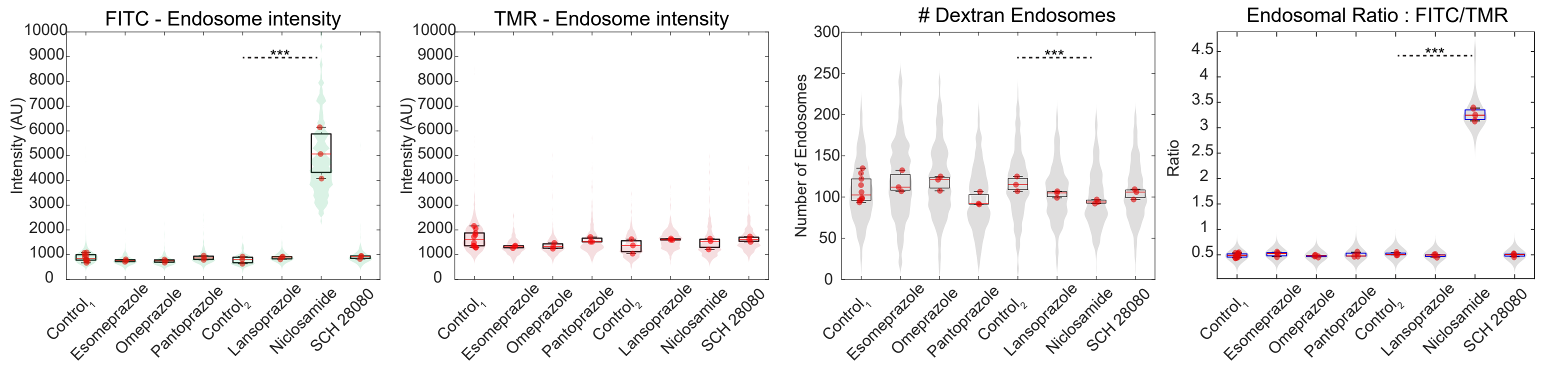

### Figure S8

#### Supplementary Figure 8

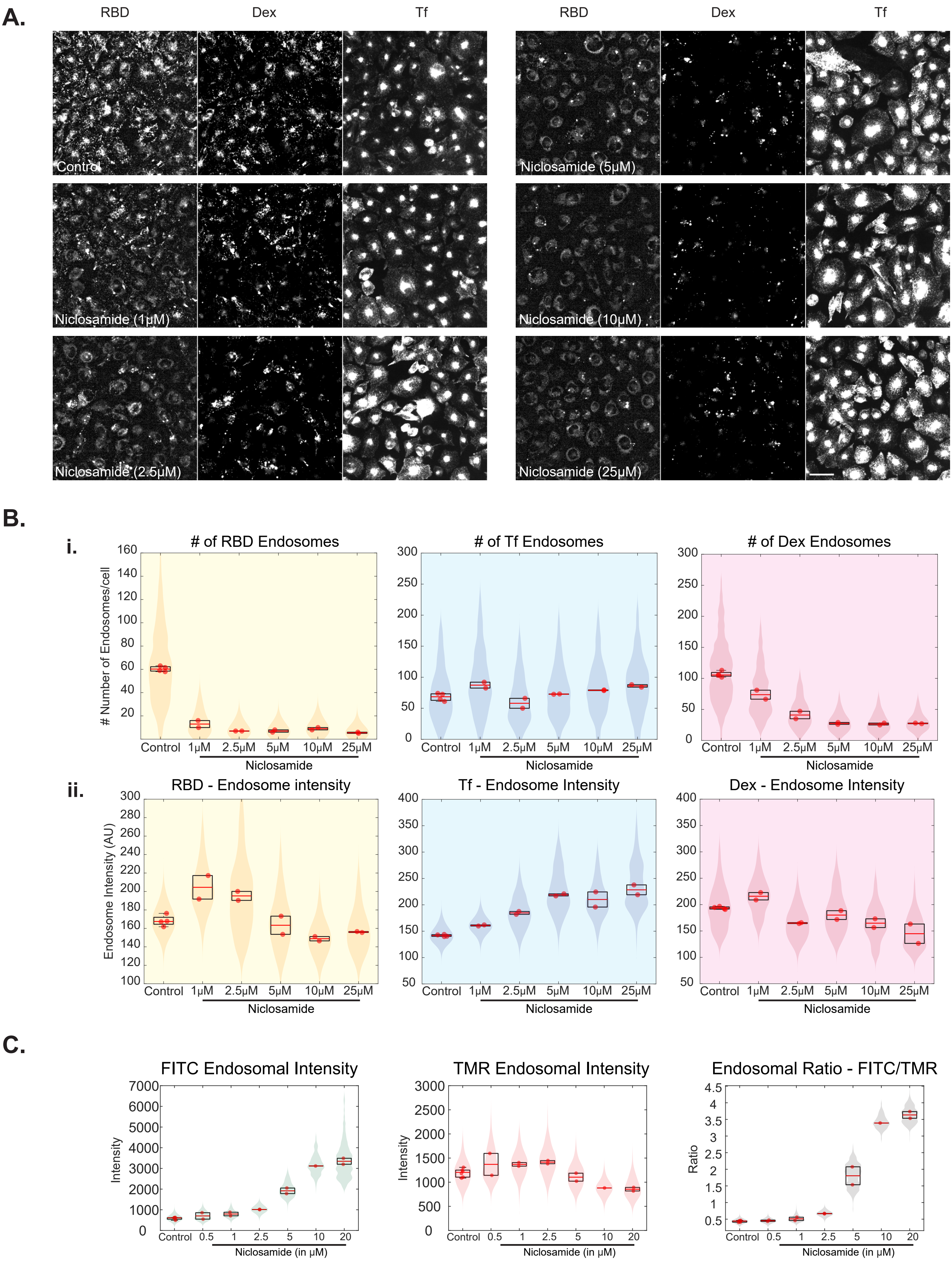

### Figure S9

#### Supplementary Figure 9

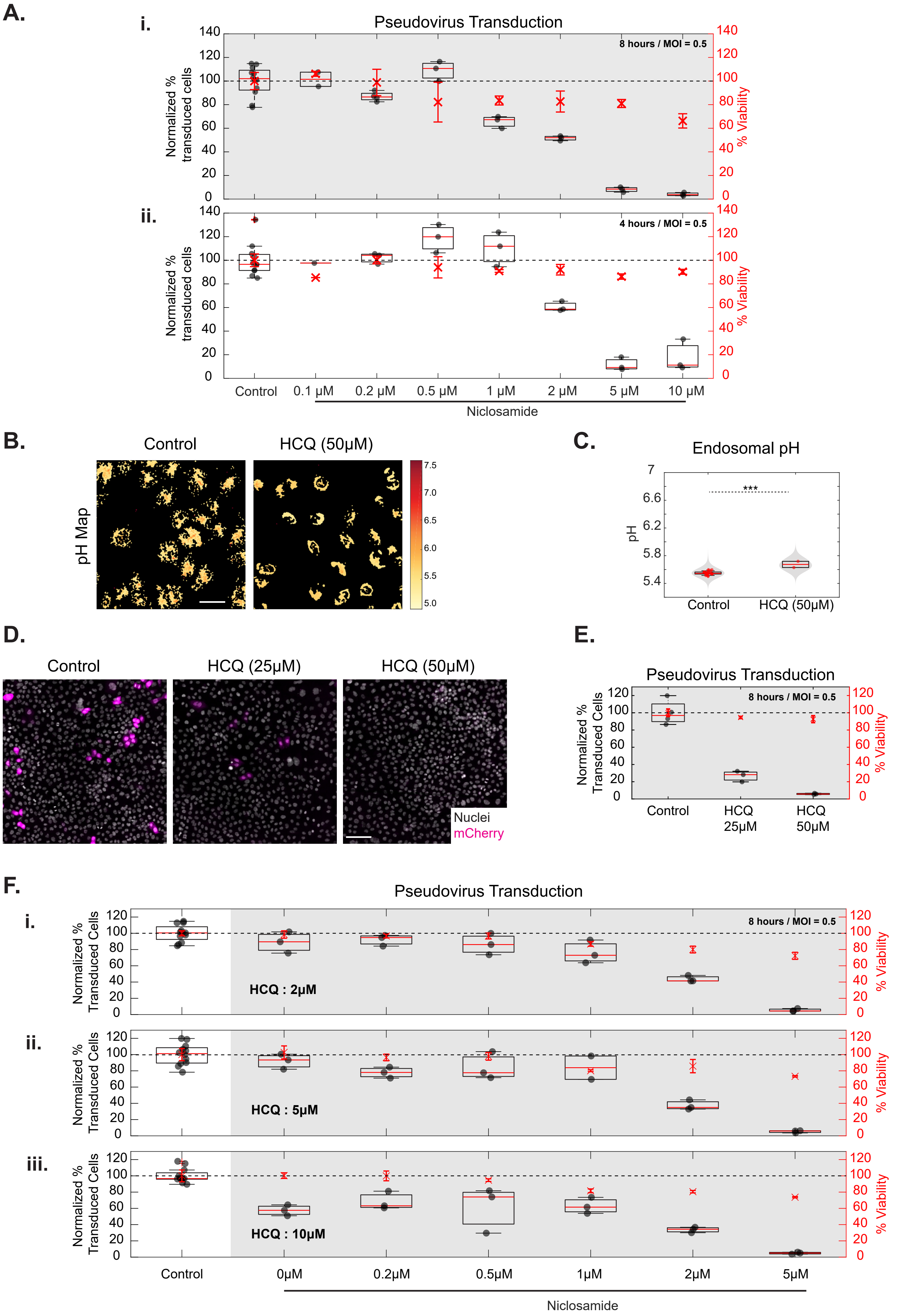

### Figure S10

#### Supplementary Figure 10

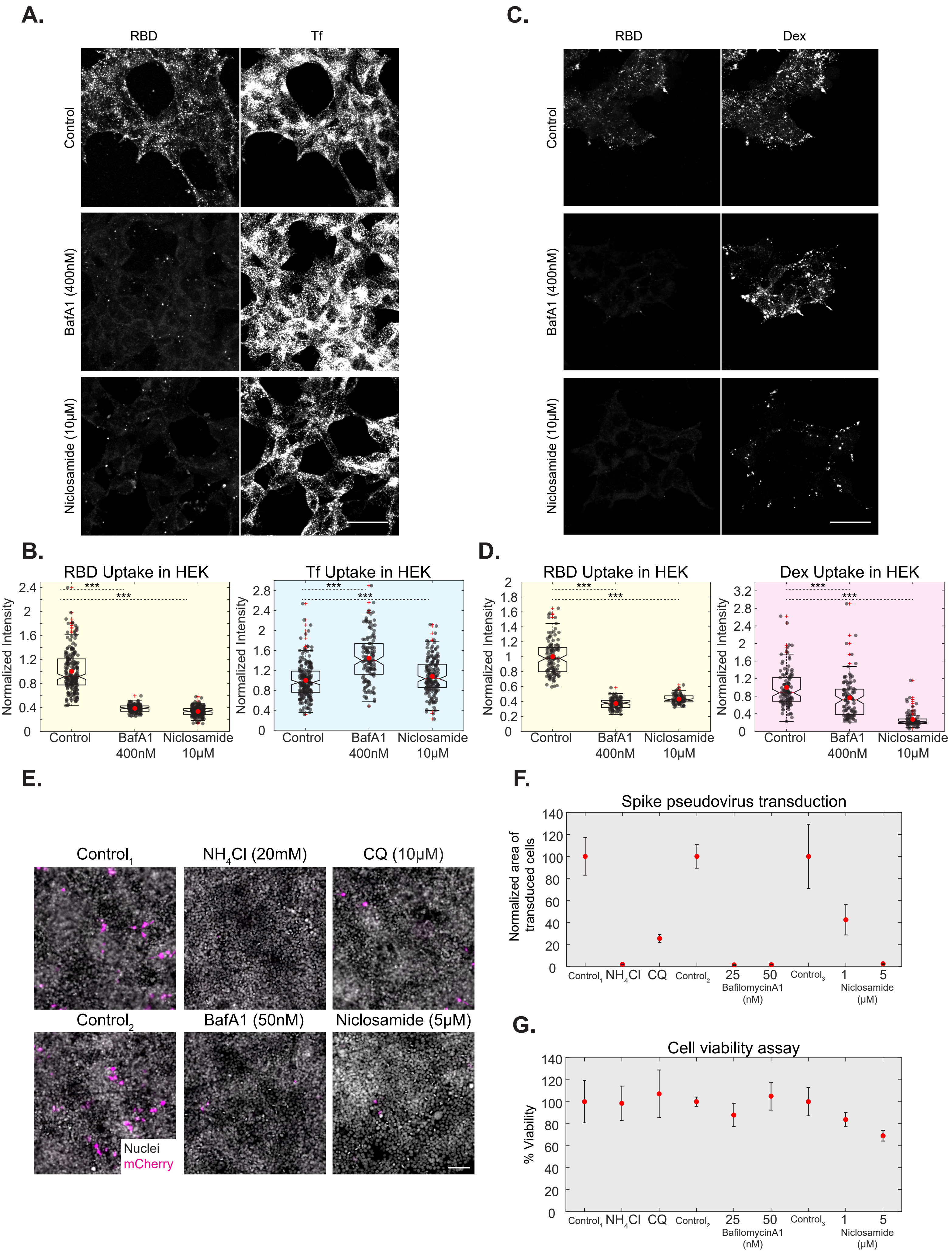
