## Supplementary Information for "Niclosamide inhibits SARS-CoV2 entry by blocking internalization through pH-dependent CLIC/GEEC endocytic pathway"

### Supplementary Figure Legends

*Supplementary Figure 1: Generation of SARS-CoV2 probe to study its endocytosis itinerary*

A: Schematic describing the protocol for purification and fluorescent labelling of RBD.

B: i) Image of a 10% SDS PAGE Gel showing the output from Ni-NTA purification of his-tagged RBD. Input is the culture supernatant containing secreted RBD (marked by a black box on the gel). FL is the flowthrough after binding the supernatant to the Ni-NTA column. RBD is eluted in fractions containing increasing concentrations of imidazole (50, 100, 150, 200, 250 mM). ii) Image of a 10% SDS PAGE Gel showing purified RBD after Gel filtration step of purification. L represents the ladder lane.

C: AGS cells were transfected with myc-ACE2 and pulsed with RBD and transferrin for 30 minutes. Surface ACE2 was marked using anti-myc antibody. Myc-ACE2 transfected cells show increased RBD.

D, E: AGS cells were transfected with myc-ACE2 and pulsed with RBD for 30 minutes. The cell surface-bound RBD was stripped using ascorbate buffer and cell surface ACE2 was labelled using anti-myc antibody. Images in D and scatter plot in E shows a positive correlation between the amount of RBD endocytosed and levels of surface ACE2. Number of cells >50.

Scale bar: 40µm (C, D).

*Supplementary Figure 2: RBD uptake is sensitive to CG pathway inhibitors*

A, B: AGS cells were pulsed with RBD, dextran and transferrin for 10 minutes and imaged at high resolution after fixation. Images in A and quantification in B shows that dextran and RBD are more correlated compared to dextran and transferrin (p-value < e-04) or transferrin and RBD (p-value < e-05) as measured using Pearson's correlation coefficient (PCC). Number of cells = 10.

C, D: AGS cells were treated with Control (0.6% DMSO) or AN96 25µM for 30 minutes, pulsed with RBD, dextran and transferrin for 30 minutes with Control or AN96 and imaged at high resolution upon fixation. Images are shown in C and quantification of Manders' co-occurrence coefficient is shown in D. This depicts the fraction of RBD endosomal intensity with transferrin or dextran (i), the fraction of transferrin endosomal intensity with dextran or RBD (ii) and the fraction of dextran endosomal intensity with transferrin or RBD (iii). As seen in D(i), in control cells, the fraction of RBD endosomal intensity is more associated with dextran than transferrin (p-value < e-07). With AN96, internalized RBD and dextran is associated more with transferrin compared to control cells.  Numbers of cells in each condition >10. Wilcoxon rank-sum test yields the following p-value table on comparing control with AN96 treated cells for different conditions:

| **D (i)** | x = Tf | x = Dex | **D(ii)** | x = Dex | x = RBD | **D(iii)** | x = Tf | x = RBD |
|---|---|---|---|---|---|---|---|---|
| % RBD | <e-04 | 0.03 | % Tf | 0.0012 | < e-04 | % Dex | <e-04 | <e-05 |

E, F: AGS cells were treated with Control or ML141 50µM for 30 minutes and pulsed with RBD and Dextran for 30 minutes with or without the inhibitor. RBD (p-value < e-9) and Dextran (p-value < e-20) uptake is significantly reduced upon treatment with ML141. Images are shown in E and quantification in F. Numbers of cells > 100 for each treatment.

G, H: AGS cells were treated with Control (0.2% DMSO) or Amiloride 1mM for 30 minutes and pulsed with RBD, transferrin and dextran for 30 minutes with or without the inhibitor. RBD (p-value = 0.05), Dextran (p-value = 0.04) and transferrin (p-value = 0.013) uptake is not altered with Amiloride. Images are shown in G and quantification in H. Numbers of cells > 80 for each treatment.

Data representation is as described in Figure 1. Scale bar: 10 µm (A, C) and 40µm (E, G).

*Supplementary Figure 3: RBD uptake is sensitive to acidification inhibitors*

A, B: AGS cells were treated with Control (0.3% DMSO, 0.6% DMSO, 0% DMSO) or inhibitors (BafA1 200nM, BafA1 400nM, $NH_4Cl$ 30mM) for 30 minutes and then pulsed with RBD, transferrin and dextran for 30 minutes with or without inhibitors. Images are shown in A and quantification in B with total cell mean intensity shown in (i), the number of endosomes shown in (ii) and intensity per endosome shown in (iii) for each probe in each condition. $Control_1$ is 0.3% DMSO, $Control_2$ is 0.6% DMSO and $Control_3$ is 0% DMSO. Number of repeats ≥ 4 for each treatment and each repeat has >80 cells.

C, D: AGS cells transfected with myc-ACE2 were treated with Control (0.2%DMSO) or BafA1 400nM or Niclosamide 10µM for 30 minutes and then pulsed with RBD for 30 minutes. The cell surface-bound RBD was stripped using ascorbate buffer and cell surface ACE2 was labelled using anti-myc antibody. Normalized RBD uptake is quantified as the ratio of the amount of internalized RBD to the amount of surface ACE2. Images depicted in C and quantification in D show that there is a reduction of RBD uptake upon treatment with BafA1 (p-value < e-08) or Niclosamide (p-value < e-07) in transfected as well as untransfected cells. Number of cells > 50 for each condition.

Data representation in B and D are as described in Figure 2 and 1, respectively. Scale bar: 40µm (A, C).

*Supplementary Figure 4: Acidification inhibitors neutralize endosomal pH*

A, B: pH calibration in AGS cells. AGS cells pulsed with pH-sensitive (FITC) and pH-insensitive (TMR) dextran were incubated in buffers of different pH with 5µg/ml of Nigericin and imaged live. A steady increase in the endosomal ratio of FITC/TMR with increasing pH is observed. The observed ratio vs clamped pH is fit to a sigmoidal curve (red curve) which is used as a calibration curve to estimate the pH of endosomes. Numbers of cells in each condition is >100 cells. The data in B is represented as mean +/- SD.

C, D: For the experiment described in Figure 2F-H, images including pH maps are shown in 2F, 2H, S4C and quantification in 2G, S4D. FITC and TMR endosomal intensities, numbers of endosomes and FITC/TMR endosomal ratio are quantified in S4D. BafA1 200nM/400nM and $NH_4Cl$ increases FITC intensity and reduces numbers of endosomes. $NH_4Cl$ also affects trafficking as seen with an increase of TMR intensity. Endosomal ratio (as a proxy for endosomal pH) also shows an increase with all the acidification inhibitors. $Control_1$ is 0.2% DMSO, $Control_2$ is 0.4% DMSO and $Control_3$ is 0% DMSO. Number of repeats ≥ 3 for each treatment and each repeat has >80 cells. Wilcoxon rank-sum test yields the following p-value table on comparing different conditions with respective controls:

| | Baf_200nM | Baf_400nM | $NH_4Cl$ |
|---|---|---|---|
| FITC Endosomal Intensity | < e-98 | < e-125 | < e-233 |
| TMR Endosomal Intensity | < e-15 | < e-34 | < e-167 |
| # Endosomes | < e-12 | < e-53 | < e-160 |
| Endosomal Ratio | < e-117 | < e-124 | < e-227 |

E, F: Estimation of FITC/TMR ratio of early endosomes. AGS cells were pulsed with FITC and TMR dextran for 20 minutes, chased for 10 minutes and imaged live. Throughout the pulse and chase duration, the cells were incubated with Control (0.2%DMSO) or BafA1 400nM or Niclosamide 10µM. Dextran uptake and TMR endosomal intensity are marginally reduced with Niclosamide while unaffected with BafA1. An increase in FITC endosomal intensity as well FITC/TMR endosomal ratio is observed with both inhibitors. Number of cells > 35 for each condition. Wilcoxon rank-sum test yields the following p-value table on comparing different conditions with respective controls:

| | BafA1 | Niclosamide |
|---|---|---|
| Dextran Uptake | 0.43 | 0.007 |
| FITC Endosomal Intensity | < e-14 | < e-13 |
| TMR Endosomal Intensity | 0.39 | < e-06 |
| Endosomal Ratio | < e-14 | < e-13 |

Data representation in D and F is as described in Figure 2 and Figure 1, respectively. Scale bar: 40µm (A, C, E).

*Supplementary Figure 5: Chloroquine does not affect RBD uptake and minimally affects endosomal acidification in AGS cells*

A, B: AGS cells were treated with Control (0% DMSO) or CQ 10µM for 30 minutes and pulsed with RBD, transferrin and dextran for 30 minutes with or without the inhibitor. Images shown in A and quantification in B show no change in the uptake of transferrin and RBD and marginal change in dextran uptake with CQ (p values = 0.2 for RBD, 0.24 for Tf, 0.013 for Dex). Box plot represents the distribution of medians of each repeat which is denoted by black dots. Number of repeats = 18 and 12 for Control and CQ, respectively and each repeat has >100 cells.

C, D: AGS cells were pulsed with FITC and TMR dextran for 2 hours, chased for 1 hour with Control (0.2% DMSO), 10, 50 or 100µM of CQ and imaged live. CQ minimally alters the FITC/TMR ratio (p-value < e-23, 0.38 and < e-03 for 10, 50 and 100 µM of CQ, respectively). Numbers of cells in each condition is >150 cells.

E, F: AGS cells pulsed with pH-sensitive (FITC) dextran for 2 hours and chased for 1 hour with Control or 100µM CQ or 15 minutes with 30mM $NH_4Cl$ and imaged live. CQ increases the 488/458 ratio of FITC dextran only slightly (p < e-10) compared to the increase brought about by $NH_4Cl$ (p-value < e-27). Images are shown in E and quantification in F. Number of cells > 75 for each treatment.

G, H: Using AGS cells treated with Control or CQ for 12 hours, RBD, dextran and transferrin uptake experiment (G) and FITC/TMR endosomal ratio estimation experiment (H) was conducted. Quantification in G and H show that RBD uptake, dextran uptake and FITC/TMR endosomal ratio are unaffected by long term treatment with CQ. Number of cells > 80 for each condition.

Data representation in B, D is as described in Figure 2, F, G and H is as described in Figure 1. Scale bar: 40µm (A, C, E).

*Supplementary Figure 6: Characterization of SARS-CoV2 spike pseudotyped virus*

A: Schematic showing the strategy for generating SARS-Cov2 Spike-pseudovirus. 2$^{nd}$ generation lentiviral helper plasmid psPAX was co-transfected with the reporter plasmid pHRmCherry and SARS-CoV2 Spike protein-encoding plasmid pTwist Spike in HEK-293T cells to generate Spike pseudotyped virus particles. pHRmCherry reporter plasmid was used to score for infected cells by mCherry expression.

B: Western blot showing bands of different molecular weights as detected by the anti-Strep-tag antibody which recognizes the C-term 2X Strep-tag on the Spike proteins incorporated into the pseudovirus particles.

C: Specificity of Spike pseudovirus was checked by infecting human origin HEK-293T cells and mouse origin NIH3T3 cells at MOI=1 for 72 hours. Quantification of percentage transduction shows that the mouse origin cell line showed significantly lower transduction than the human cell line. Number of repeats = 2 for each condition. The data is plotted as mean +/- SD.

D: Specificity of Spike protein-ACE-2 dependent pseudovirus entry was tested in a competition assay in the presence of excess purified RBD of Spike. Quantification of percentage transduction shows a reduction in transduction efficiency with Spike RBD in HEK-293T cells. Number of repeats = 2 for each condition. The data is plotted as mean +/- SD.

E: Characterization of transduction efficiency in AGS cells of Spike-pseudovirus at varying MOIs and varying incubation times. Quantification of percentage transduced mCherry positive cells depicts a steady increase in transduction as a function of MOI and incubation times. Number of repeats = 2 for each condition. The data is plotted as mean +/- SD.

F, G: Images of AGS cells expressing the reporter mCherry protein upon transduction with Spike-pseudovirus in F and quantification in G show no significant reduction in transduction efficiency upon treatment with 5µM of ML141 compared to control (p-value = 0.55). Number of repeats = 3,2 for control (0% DMSO) and ML141, respectively.

Data representation in G is as described in Figure 3. Scale bar: 40µm (F).

*Supplementary Figure 7: Identifying FDA-approved drugs functioning similar to BafA1 and $NH_4Cl$*

A: For the experiment described in Figure 4B–4C, images are shown in 4B and S7A and quantification in 4C. Wilcoxon rank-sum test yields the following p-value table on comparing the distribution of different inhibitors with respective controls for the uptake of RBD, transferrin and dextran:

| Uptake | Esomeprazole | Omeprazole | Pantoprazole | Lansoprazole | Niclosamide | SCH 28080 |
|---|---|---|---|---|---|---|
| RBD | < e-06 | < e-14 | 0.003 | < e-05 | < e-195 | < e-12 |
| Tf | 0.013 | < e-34 | < e-38 | < e-04 | < e-155 | < e-20 |
| Dex | 0.005 | 0.113 | 0.011 | 0.592 | < e-133 | < e-06 |

B, C: For the experiment described in Figure 4D–4E, images including pH maps are shown in 4D and S7B and quantification in 4E and S7C. FITC and TMR endosomal intensities and numbers of endosomes are quantified in S7C. Niclosamide increases FITC intensity, reduces numbers of endosomes and has minimal effect on TMR intensity. Wilcoxon rank-sum test yields the following p-value table on comparing the distribution of different inhibitors with respective controls for the listed properties:

| | Esomeprazole | Omeprazole | Pantoprazole | Lansoprazole | Niclosamide | SCH 28080 |
|---|---|---|---|---|---|---|
| FITC Endosomal Intensity | < e-35 | < e-46 | < e-06 | < e-14 | < e-112 | < e-14 |
| TMR Endosomal Intensity | < e-53 | < e-49 | < e-14 | < e-25 | 0.003 | < e-21 |
| Numbers of Endosomes | < e-05 | < e-05 | < e-04 | < e-03 | < e-07 | 0.004 |
| Endosomal Ratio | < e-18 | 0.16 | < e-03 | < e-15 | < e-110 | < e-09 |
| pH | < e-19 | 0.18 | < e-03 | < e-15 | < e-110 | < e-10 |

*Supplementary Figure 8: Niclosamide functions like BafA1 in inhibiting RBD uptake and neutralizing the endosomal pH*

A, B: For the endocytic assay experiment described in Figure 5A, images are shown in S8A and quantification in 5A and S8B, with total cell mean intensity shown in 5A, number of endosomes shown in S8B(i) and intensity per-endosome shown in S8B(ii). The number of RBD endosomes and dextran endosomes decrease, while transferrin endosomal intensity increases with increasing concentrations of Niclosamide. Wilcoxon rank-sum test yields the following p-value table on comparing the distribution of uptake of the indicated concentrations of Niclosamide with control:

| Niclosamide in µM | 1 | 2.5 | 5 | 10 | 25 |
|---|---|---|---|---|---|
| RBD Uptake | < e-126 | < e-181 | < e-221 | < e-199 | < e-234 |

| | | | | | |
|---|---|---|---|---|---|
| Tf Uptake | < e-34 | < e-104 | < e-151 | < e-140 | < e-158 |
| Dex Uptake | < e-09 | < e-51 | < e-124 | < e-138 | < e-179 |

C: For the pH estimation assay described in Figure 5B–5C, quantification of endosomal FITC intensities, TMR intensities and FITC/TMR endosomal ratio is shown in S8C. A dose-dependent increase in FITC endosomal intensity, as well as ratio, is seen with increasing Niclosamide concentrations. Wilcoxon rank-sum test yields the following p-value table on comparing the distributions of indicated concentrations of Niclosamide with the control for the listed properties:

| Niclosamide (in µM) | 0.5 | 1 | 2.5 | 5 | 10 | 20 |
|---|---|---|---|---|---|---|
| Endosomal Ratio | < e-201 | < e-134 | < e-129 | < e-147 | < e-143 | < e-25 |
| pH | < e-06 | < e-72 | < e-252 | < e-245 | < e-126 | < e-203 |

Data representation in B, C is as described in Figure 2. Scale bar shown in A is 40µm.

*Supplementary Figure 9: Niclosamide and Hydroxychloroquine affect Spike-pseudovirus transduction in a dose-dependent manner*

A: Quantification in A shows the normalized percentage transduction of Spike-pseudo virus across different concentrations of Niclosamide for incubation times of 8hours(i) and 4hours(ii). Both the inhibitor and the virus were removed beyond the indicated times and cells were incubated with the continued presence of 100nM Niclosamide or 0.005% DMSO until termination. Images shown in 5D and dose-response curve depicted in 5E are related to the experiment in S9A(i). Number of repeats = 3 for each concentration of Niclosamide except 2 and 4 for 0.1µM and 0.2µM, respectively for the 8 hours set and 1 for 0.1µM Niclosamide in the 4 hours set. Wilcoxon rank-sum test yields the following p-value table on comparing the control with different concentrations of Niclosamide for different timepoints of viral incubation:

| Niclosamide (µM) | 0.1 | 0.2 | 0.5 | 1 | 2 | 5 | 10 |
|---|---|---|---|---|---|---|---|
| 4 hours | 0.52 | 0.99 | < e-04 | 0.27 | < e-99 | < e-21 | < e-103 |
| 8 hours | 0.48 | < e-04 | 0.02 | < e-23 | < e-133 | < e-46 | < e-126 |

B, C: AGS cells pulsed with pH-sensitive (FITC) dextran for 2 hours and chased for 1 hour with Control or 50µM HCQ and imaged live. HCQ increases pH only slightly. pH maps are shown in B and quantification in C. Numbers of repeats: Control = 22, HCQ = 2.

D, E: Images of AGS cells expressing the reporter mCherry protein upon transduction with Spike-pseudovirus in D and quantification in E show a dose-dependent reduction in transduction efficiency upon treatment with HCQ at the two concentrations tested compared to control (p-value < e-66 for HCQ 25µM, p-value < e-96 for HCQ 50µM). Number of repeats = 4 for control (0% DMSO) and 3 each for each concentration of HCQ.

F: For the experiment described in Figure 5F–5G, Quantification in S9F shows the normalized percentage transduction across indicated concentrations of Niclosamide in combination with indicated Hydroxychloroquine concentration of 2µM(i), 5µM(ii) and 10µM(iii). The percentage of cell viability for each condition is also indicated. Number of repeats = 2 for HCQ 5µM + Niclosamide 1µM combination and 3 each for all other combinations. Wilcoxon rank-sum test yields the following p-value table on comparing different inhibitor conditions with respective controls:

| Concentration in µM | Nic_0 | Nic_0.2 | Nic_0.5 | Nic_1 | Nic_2 | Nic_5 |
|---|---|---|---|---|---|---|
| HCQ_2 | 0.0025 | 0.031 | < e-04 | < e-11 | < e-61 | < e-129 |

| | | | | | | |
|---|---|---|---|---|---|---|
| HCQ_5 | 0.15 | < e-05 | < e-03 | < e-03 | < e-48 | < e-110 |
| HCQ_10 | < e-34 | < e-19 | < e-06 | < e-28 | < e-77 | < e-124 |

Data representation in A, E and F are as described in Figure 3 and C as described in Figure 2. Scale bar: 100µm (B).

*Supplementary Figure 10: BafA1 and Niclosamide affect RBD uptake and reduce Spike pseudovirus infection in HEK-293T cells*

A-D: HEK-293T cells were treated with Control (0.4%DMSO), BafA1 400nM or Niclosamide 10µM for 30 minutes and pulsed with RBD and transferrin (A and B) or RBD and dextran (C and D) for 30 minutes with or without inhibitors. Images are shown in A and C, quantification is shown in B and D. RBD and dextran uptake is robustly reduced, while transferrin uptake increases upon treatment with BafA1 and Niclosamide. Number of cells ≥ 75 for each treatment. Wilcoxon rank-sum test yields the following p values on comparing the uptake different inhibitors with respective controls:

| **(B)** | BafA1 | Niclosamide | **(D)** | BafA1 | Niclosamide |
|---|---|---|---|---|---|
| RBD | < e-53 | < e-60 | RBD | < e-34 | < e-30 |
| Tf | < e-17 | 0.0097 | Dex | < e-04 | < e-26 |

E: Images show Spike-pseudovirus transduced mCherry positive HEK-293T cells in the presence of $NH_4Cl$ 20mM, CQ 10µM, BafA1 50nM and Niclosamide 5µM.

F: Quantification of the normalized area of mCherry positive cells as a proxy for transduction, shows a reduction in transduction efficiency upon treatment with $NH_4Cl$ 20mM and CQ 10µM compared to 0% DMSO ($Control_1$), 25nM and 50nM of BafA1 compared with 0.05%DMSO ($Control_2$), 1µM and 5µM of Niclosamide compared to 0.1%DMSO ($Control_3$). Number of repeats = 3 for each condition except 4 for 0%DMSO. The data is represented as mean +/- SD.

G: Toxicity, as assessed by MTT based colorimetric assay, is represented as percentage viability of cells upon treatment with $NH_4Cl$ 20mM and CQ 10µM compared to 0% DMSO ($Control_1$), 25nM and 50nM of BafA1 compared with 0.05%DMSO ($Control_2$), 1µM and 5µM of Niclosamide compared to 0.1%DMSO ($Control_3$). Number of repeats = 3 for each condition except 9 for 0%DMSO. The data is represented as mean +/- SD.

Data representation in B and D are as described in Figure 1. Scale bar: 40µm (A, C), 100µm (E).

### Supplementary Methods:

*Cell lines, constructs, and antibodies:*

Cell lines: AGS cells (Human adeno gastric carcinoma cells stably expressing human Folate Receptor), HEK 293T and NIH3T3. AGS cells were maintained in HF12 media (HiMEDIA, India). HEK293T and NIH3T3 cells were maintained in DMEM-High Glucose (HiMEDIA, India). All culture media were supplemented with 10% FBS (16000044, Invitrogen), $NaHCO_3$ and L-Glutamine/ Penicillin/ Streptomycin solution (G1146, Sigma Aldrich). For AGS cells, the antibiotic hygromycin B (10687010, Invitrogen) was added at a concentration of 200µg/ml for selection during maintenance of cultures and the antibiotic-free medium was used for assays.

Constructs: pCAGGS-Secreted-RBD-6X-His (gift from Florian Krammer, Mt. Sinai), pCEP4-myc-ACE2 (gift from Erik Procko; Addgene plasmid #141185), pTwist-EF1alpha-nCoV-2019-S-2xStrep (gift from Nevan Krogan, UCSF), pHR mCherry plasmid (gift from Minhaj Sirajuddin, inSTEM, India) and second-generation lentiviral helper plasmid psPAX2 (gift from Didier Trono; Addgene plasmid #12260).

Antibodies: Anti-human transferrin receptor antibody was purified from mouse hybridoma and labelled using NHS-ester chemistry. Anti-Myc Mouse mAb from CST (9B11 clone; Cat# 2276) was used to label myc-ACE2. Alexa Fluor 568 labelled goat anti-mouse secondary (115-005-071; Jackson Laboratory) was used to detect labelled Myc. Monoclonal Anti-Strep Tag antibody (clone GT517; Cat# SAB2702215) from Sigma-Aldrich and Goat anti-Mouse HRP secondary antibody (115-035-208; Jackson Laboratory) was used to identify the C-term Strep-tag on the Spike protein in the Spike-pseudotyped virus lysate western blot.

*Chemicals and reagents:*

BafilomycinA1 (B1793 Sigma), $NH_4Cl$ (A9434, Sigma), ML141 (4266, Tocris Bioscience), Chloroquine diphosphate (C6628, Sigma), Amiloride (A7410, Sigma), Atto 488 NHS Ester (41698; Sigma), TMR dextran (D1817, Thermo Scientific), Dextran 10kDa (D1860, Invitrogen), FITC Isomer-I (F1906, Molecular Probes), MTT (3-(4,5-Dimethylthiazol-2-yl)-2,5-diphenyltetrazolium bromide, M6494, Invitrogen), Hoechst (bisBenzimide trihydrochloride, H33342, Merck), Fugene 6 Transfection Reagent (E2692, Promega), Lipofectamine 3000 (Invitrogen), Lenti-X concentrator (631232, Takara Bio), 10X lysis buffer (9803, CST), Micro BCA™ Protein Assay Kit (23235; Thermo Scientific), PVDF membrane (Immobilon-P, IPVH00010, Millipore) and SuperSignal™ West Pico Chemiluminescent Substrate (34080, Thermo Scientific). Hydroxychloroquine was provided by Mylan Laboratories.

*Preparation of RBD:*

RBD purification was optimized using a protocol from Krammer's Laboratory [1]. The procedure is briefly described in Figure S1A. HEK-293T cells were seeded in T75 flasks, grown up to 60% confluency and transfected with secreted RBD construct using Lipofectamine 3000. Cells were incubated with serum-free media during the first 4 hours of transfection and then changed to medium containing 2% serum. Cells were grown under these conditions for 40-48 hours, after which the media with secreted RBD was collected. The same cells were further incubated with fresh 2% serum-containing media for another 40-48 hours and the media containing secreted RBD was collected at the end of the incubation time. The collected media was filtered using a 0.45-micron filter to remove cell debris and other contaminants. RBD was purified by immobilized metal ion affinity chromatography

(IMAC) over a 5-mL HiTrap His column (GE Healthcare Life Sciences) with step elution using buffers containing 0-500mM Imidazole. RBD containing fractions was found to be eluted within the range of 150 - 300mM Imidazole-containing elution buffers (Figure S1B). These fractions containing RBD were pooled and concentrated using 3kDa filters (Amicon). The concentrated filtrate was loaded onto HiLoad® 16/600 Superdex® 200 (GE Healthcare Life Sciences) gel filtration column and eluted using 1X PBS to separate RBD from serum albumin. The RBD-containing fractions were collected and concentrated using Centrivap vacuum concentrator or 3 kDa filters (Amicon). To label RBD using NHS ester chemistry, a 10-fold molar excess of Atto-488 NHS ester (Sigma) was incubated with RBD at pH 8.5 for an hour at room temperature. Labelled protein was separated from free dye using gravity flow-based size exclusion chromatography (Thermo Scientific).

*Preparation of Spike-pseudovirus particles and estimation of titre:*

Spike-pseudovirus particles were generated in HEK-293T cells by co-transfecting the lentiviral helper plasmid psPAX, the C-term 2X-Strep-tagged Spike-encoding pTwist plasmid and a lentiviral pHR mCherry reporter plasmid encoding soluble mCherry protein under the control of SFFV promoter (Figure S6A). Transfection was carried out using Lipofectamine 3000. The supernatants were collected at least 3 times post-transfection between 48hours-72 hours. The viral supernatants were pooled and concentrated using the Lenti-X concentrator following manufacturer's protocol. The concentrated virus was resuspended in a small volume of Opti-MEM® (Gibco®), aliquoted and stored in -80ºC. For determining titre, AGS cells were transduced at different dilutions of the pseudovirus preparation for 72hours. The number of mCherry positive colonies in the entire area of a 96 well plate was counted and titre was recorded as colony-forming units/ml (CFUs/ml). All procedures were carried in containment in a BSL2 facility.

*Western blotting:*

Virus-containing culture media, from 48-72h post-transfection, were pooled as mentioned in the above section. The medium was briefly centrifuged to remove cell debris and filtered using a 0.45µm filter. Filtered supernatant was overlaid on freshly prepared sucrose solution layer (50mM Tris-HCl, pH 7.4, 100mM NaCl and 0.5mM EDTA; with 20% sucrose (w/w) at a 4:1 ratio (v/v) [2]) and centrifuged at 15000 g at 4ºC for 4h. The supernatant was carefully discarded and any remaining liquid was removed by gently inverting the tube onto a blotting paper. The pellet containing pseudovirus particles was gently resuspended in a minimum volume of PBS (pH 7.4) on ice. Pseudovirions were then lysed using 10X lysis buffer. Protein content was estimated using a Micro BCA™ Protein Assay Kit as per manufacturer's protocol. To validate the presence of spike protein, 20µg of this lysate was loaded onto a 10% SDS polyacrylamide gel, blotted on a PVDF membrane and probed with anti-Strep-tag primary antibody followed by HRP secondary antibody.

*RBD competition assay:*

HEK-293T cells were pre-incubated in medium containing excess purified RBD for 10 minutes at 37°C. An excess of $10^{12}$ molecules of RBD was used for 1500 pseudovirus particles. Pseudovirus particles are estimated to have $4.5*10^5$ binding sites, considering 100 spike trimer or 300 spike monomers per virus particle, using the reported numbers for SARS-CoV1 [3]. Following this, pseudoviruses were added to cells in the continued presence of excess RBD for 4 hours. After this, the medium was removed, and cells were replenished with fresh medium (not containing RBD or pseudoviruses) for 60 hours. Numbers of mCherry-positive cells was estimated and reported in Figure S6D.

*Cell viability assay:*

To check the toxicity of inhibitors in HEK-293Ts, cells were treated with specified concentrations of the inhibitors, similar to the format employed for transduction assays (except no pseudoviruses were added). At the end of the incubation period, cells were washed twice with serum-free medium and then incubated with MTT working solution (20µl of 5mg/ml MTT in serum-free medium + 80µl of growth medium) for 3 hours at 37°C. Following this, contents from all wells were removed completely and 100µl of DMSO was added to each well. Complete dissolution of crystals in DMSO was ensured. Optical densities were then recorded at 570nm. Wells containing no cells were used for background correction. Values corresponding to cells treated with respective vehicle controls were considered as 100%.

*High content screening and automated imaging methodologies:*

For high-throughput endocytic and pH estimation assays, AGS cells were plated in optical bottom 96 or 384 well plates at ~8000 or ~2000 cells/well and processed 16 hours after seeding. Experiments were conducted as detailed in the main text Methods using an automated plate washer and a robotic arm.

*Image segmentation and feature extraction:*

The maximum projected images were corrected for the system background and uneven illumination. The background was measured from wells not containing cells and subtracted from the raw data. For each channel, the uneven excitation was estimated by averaging all the maximum projected images from a single day's experiment and applying a 2D Gaussian filter of 50. This filtering removed any potential low-intensity features. The average of all the pixel values was then normalised to 1 to generate an illumination profile image. All background-corrected images were divided by this estimated value of illumination.

For cell segmentation, the Hoechst-stained nuclei images were used to first identify cells using Otsu's method on log-scaled intensity images. Next, using Sobel edge detection on each endosome channel, a 2D Gaussian filter of 20 pixels together with the Otsu methods allowed for the generation of a binary cell mask that contained most of the cell body of each cell. This allowed us to determine a field averaged background value by measuring the median intensity of the pixels outside of the area occupied by cells for all channels. Finally, individual endosomes within a cell were identified by using the endosome channels together with Otsu's method using the expected endosome sizes between 1-15 pixel diameters. Having identified cells and individual endosomes, their mean intensities in all channels, as well as their size and shape, were determined.

*Synthesis of Niclosamide:*

Niclosamide was synthesized by coupling 5-chlorosalicylic acid with 2-chloro-4-nitroaniline in the presence of thionyl chloride.

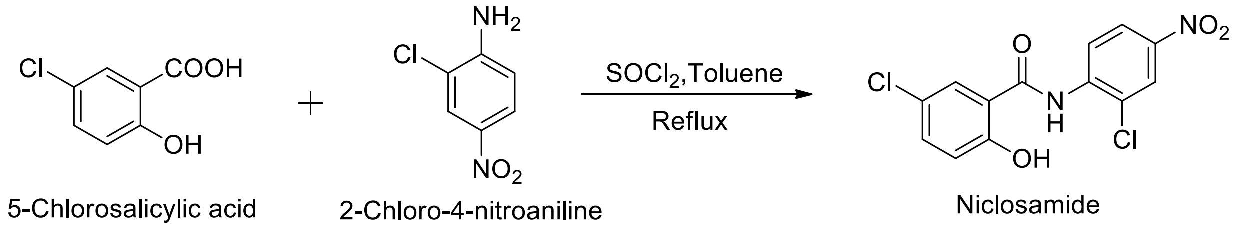

The 5-chlorosalicylic acid (1.72 g, 10 mmol) was suspended in 20 mL toluene. Thionyl chloride (160 µL, 2.2 mmol) was added and the reaction mixture was stirred at 110 °C for two hours. To this mixture was added 2-chloro-4-nitroaniline (1.14 g, 6.6 mmol) and the reaction was continued to stir at 110 °C for another 6 hours. The reaction mixture was allowed to cool to room temperature and kept overnight without disturbing. Niclosamide gets precipitated at the bottom of the RBF. The supernatant was decanted, the precipitate was washed with an excess quantity of water and the product was dried to get 1.8 g (83% yield) of niclosamide. The product was characterized by comparison of the spectral data with literature values [4]. Light yellow powder; m.p. 225-228 °C; HPLC: 98.9% purity; $^{1}H$ NMR (400 MHz, DMSO-$d_6$) δ 12.48 (s, 1H, OH), 11.34 (s, 1H, NH), 8.80 (d, $J$ = 9.2 Hz, 1H), 8.44 (d, $J$ = 1.6 Hz, 1H), 8.30 (dd, $J$ = 8.4, 2.0 Hz, 1H), 7.98 (d, $J$ = 2.0 Hz, 1H), 7.55 (dd, $J$ = 8.2, 2.0 Hz, 1H), 7.10 (d, $J$ = 8.8 Hz, 1H); $^{13}C$ NMR (100 MHz, DMSO-$d_6$) δ 162.56, 155.18, 142.54, 141.17, 134.03, 130.07, 124.78, 123.90, 123.78, 122.37, 120.70, 119.41, 119.16; IR ($\nu_{max}$): 3488, 2921, 1679, 1607, 1348, 1195, 1124, 901 $cm^{-1}$; ESI-MS: *m/z* 325.05 $[M-H]^-$ .

Spectral scans of Niclosamide:

**$^{1}$H NMR:**

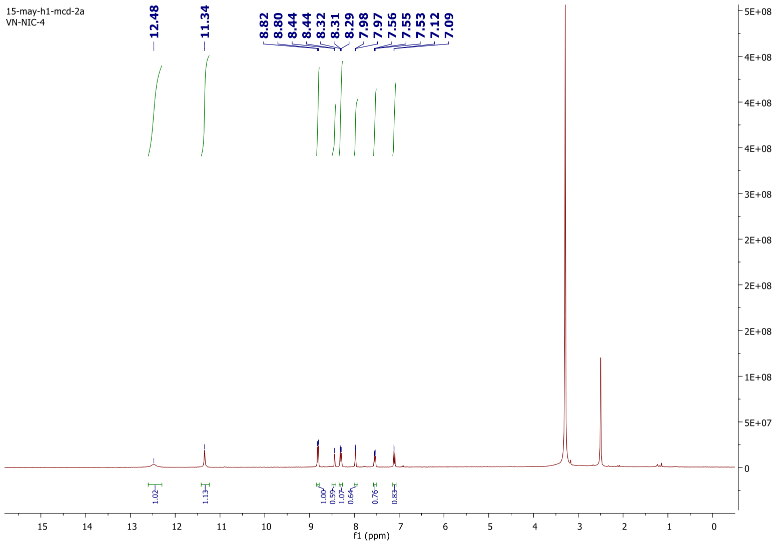


**$^{13}$C NMR:**

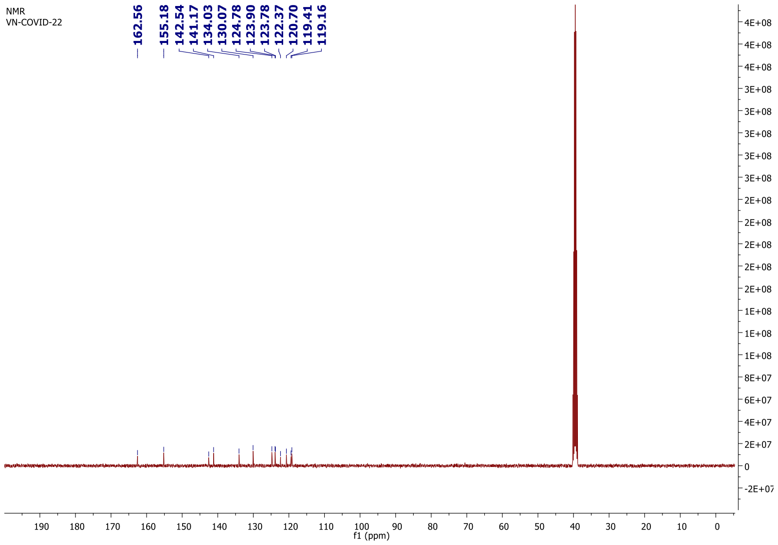

*AN96, a stable analog of GBF1 inhibitor, LG186:*

AN96 (4-(2-(3,4,5-trimethoxybenzylidene)hydrazineyl)-5,6,7,8,9,10-hexahydrocycloocta[4,5]thieno[2,3-d]pyrimidine) was synthesized to create a more stable of analog of LG186[5,6], and this will be detailed elsewhere (Godbole et al., Manuscript in preparation).

**LG-186** [5,6]

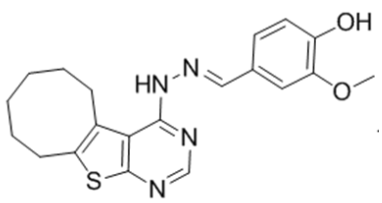

**AN96**

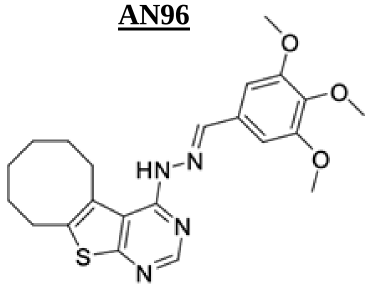

*Extraction of Esomeprazole and Pantoprazole:*

The commercially available tablets were dissolved in the aqueous phase and extracted with ethyl acetate (3x50mL). The organic layer was passed through anhydrous sodium sulphate, and the solvent was evaporated using Rotavapor. Recrystallization with hexane gave the pure compound in powder form. The purity of the drug has been confirmed using $^1$H-NMR spectroscopy. Molecular weight of Esomeprazole and Pantoprazole is 345.4 g/mol and 383.4 g/mol, respectively.

**Pantoprazole**

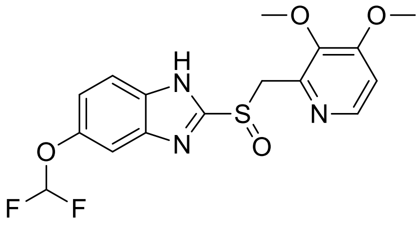

$^1$H NMR (600 MHz, DMSO-d6), δ (ppm) = 3.89 (s, 6H), 4.32 (d, J=12.9 Hz, 2H), 6.74 (dd, J=2.4 Hz, 2H),7.018 (s,1H), 7.07 (d, J=5.52 Hz, 2H), 7.24 (d, J=2.34 Hz, 1H), 7.45 (d, J=8.58 Hz, 1H).

$^{13}$C NMR (150 MHz, DMSO-d6), δ (ppm) = 56.36, 56.41, 61.42, 107.91, 108.32, 111.73, 116.26, 118.11, 119.65, 144.72, 144.80, 145.03, 146.27, 147.59.

**Esomeprazole**

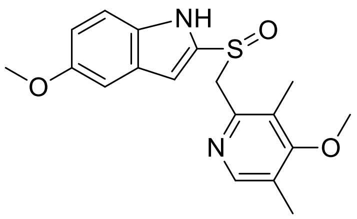

$^1$H NMR (600 MHz, DMSO-d6), δ (ppm) =2.21 (s, 6H), 3.68 (s, 6H), 4.40 (s, 2H),7.026 (s, 1H), 7.37 (d, J= 8.8 Hz, 1H), 8.25 (s, 1H).
